# A closer look at Type I left-handed β-helices provides a better understanding in their sequence-structure relationship: towards their rational design

**DOI:** 10.1101/2023.06.27.546660

**Authors:** Maxime Naudé, Peter Faller, Vincent Lebrun

**Affiliations:** Institut de Chimie (UMR 7177), University of Strasbourg – CNRS, 4 Rue Blaise Pascal, 67081 Strasbourg, France

**Keywords:** repeat protein, β-solenoid, sequence-structure relationship, protein design

## Abstract

Understanding the sequence-structure relationship in protein is of fundamental interest, but has practical applications such as the rational design of peptides and proteins. This relationship in the Type I left-handed β–helix containing proteins is updated and revisited in this study. Analysing the available structures in the Protein Data Base, we could describe further in details the structural features that are important for the stability of this fold, as well as its nucleation and termination. This study is meant to complete previous work, as it provides a separate analysis of the N-terminal and C-terminal rungs of the helix. Particular sequence motifs of these rungs are described along with the structural element they form.

## Introduction

It is commonly accepted that protein’s function arises from their structure, which itself is encoded in the sequence according to the Anfinsen’s principle.^1^ The sequence of proteins with tandem repeats is composed of a repetition of a short sequence, usually forming an elementary structural unit.^2–4^ They are particularly suitable for protein design, with their interactions it allow to conceive objects that can be tunable, leading to interesting properties regarding the design of repetitive units that are reminiscent of legos.^3,5–7^ Some databases dedicated to Structural Repeats in Proteins have been developed to facilitate their identification and analysis.^4,8–10^

Noteworthy, such protein folds can be used as domains to mediate oligomerization of chosen protein in a predictable way,^11–14^ allowing the design of higher order assemblies for example.^15–18^ So far, most designed repeat proteins were based on α-helical units (eg AlphaRep, Ankyrin repeat).^7,19,20^ Besides, β-solenoids are particularly represented in the fiber-forming proteins, such as the cytoskeletal bactofilin BacA for examples, and hence have been used to design nano-wires for example.^2,21–24^ β-solenoids are defined as a single polypeptide backbone winding around an axis, forming β-sheets and β-turns.^2,3,25^ The β-solenoids are appealing because of their mechanical properties and stability, as the amino acids oriented inwards form a hydrophobic core that stabilize the folding.^26,27^. This super structure was first discovered by Yoder et al. in 1993.28 Different types exist, depending on the number of β-strands per rung, on the number of amino acids (a.a) per rung, etc.^2,25^ For example, the β–helix designates a β-solenoid with three β-sheets per rung (triangular section), and can be right-handed or left-handed. This latter, is subdivided in two types. The Type II left-handed β–helix (eg antifreeze protein CfAFP) is composed of 15 amino acids per rung versus 18 amino acids per rung for the Type I left-handed β–helix (eg acetyltransferase LpxA or γ– carbonic anhydrase). Another difference is that the Type I is usually a trimerization domain (Figure 1a), while the Type II remains monomeric.^29^

**Figure 1.**
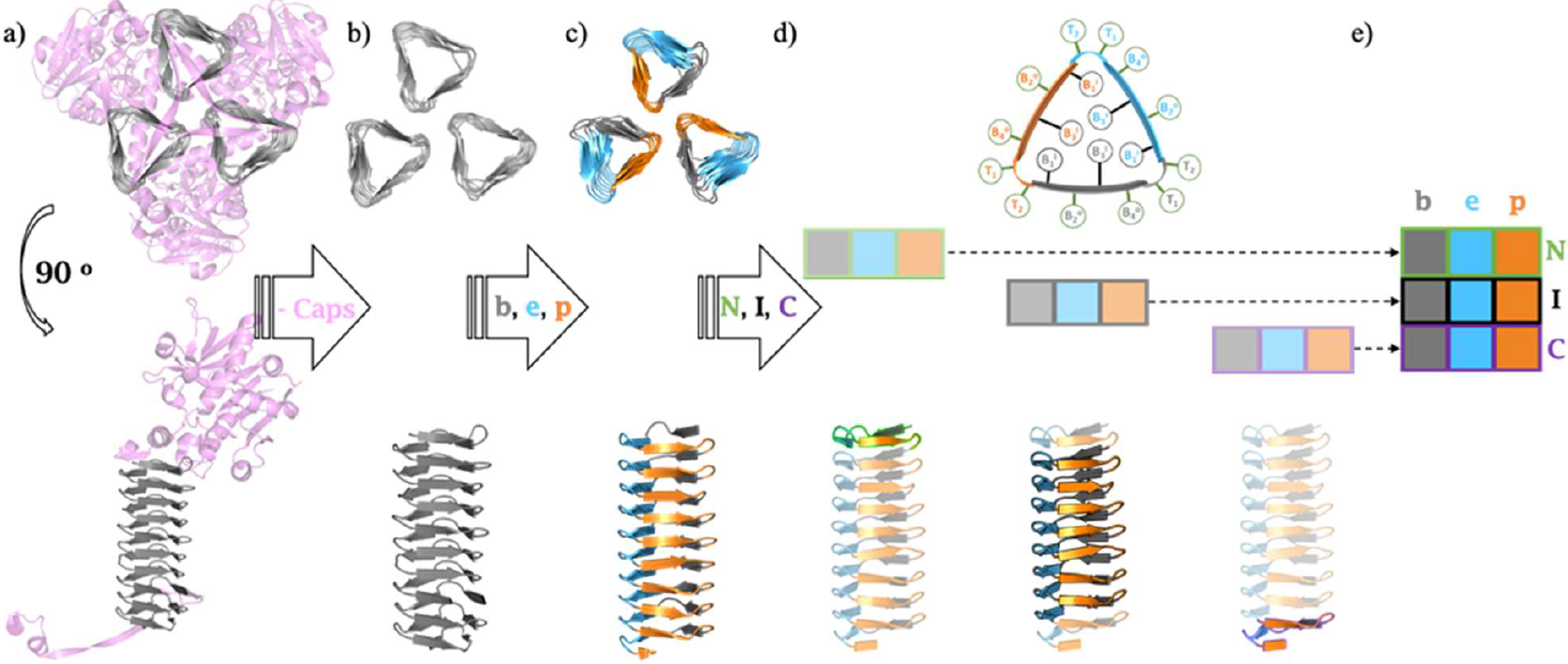
Methodology following for this LβH-I analysis. a) Selection of trimeric LβH-I crystallographic structures from the PDB and identification of the canonical folding. b) Removal of structural elements that to not belong to the canonical LβH-I. c) Labelling of the faces of the LβH-I as a function of their exposition to the solvent (buried (b), exposed (e), and partial (p)) originating from the trimeric arrangement. d) Separation of the different types or rungs: N-terminal (N), inner (I) and C-terminal (C). e) The analysis of these nine different hexads finally yields a table reporting the occurrence of the amino-acids in the 54 different positions identified.

The first structural study on Type I left-handed β–helix (hereafter abbreviated as LβH-I)domain, based on seven LβH-I protein structures, mostly focused on the torsion angles within the main chain.^30^ They distinguished the β sheets of the helix in its trimeric form, and showed that the LβH-I sequence can be built from a repetition of a hexa-peptide motif. They highlighted a few important features in the sequence, such as the importance of glycine residues at the fourth position of the β strands, the need of small residue at the first position, and the composition of the hydrophobic core.

Later, Cohen and coworkers (i) confirmed that the sequence of a LβH-I is essentially the repeat of a hexad, (ii) introduced the nomenclature for it, ie two residues for the turn (called “loop” before) (T1 and T2) and four residues for the β–strand (B1 to B4), and (iii) reported the position-dependent residue propensities. Of note, the latter was performed based on the eight crystal structures available at that time (by sequence alignment), and excluding the rungs at the N-ter and C-ter extremities.^29^ This work was completed with an experimental study assessing the tolerance of the LβH-I fold for the different amino-acids at each position of the hexad.31 In parallel, modelling studies also contributed, for example in evidencing the importance of proline and glycine residues in LβH-I for maintaining the integrity of the fold.^32^

While these previous studies established the first understanding of the sequence-structure relationship in LβH-I (compatibility and occurrence of residues in the hexad) they suffer from two limitations. Indeed, the reduction of the whole LβH-I sequence to only one hexad means considering (i) all three sides of the helix as equivalents, while they are not exposed to the same environment (due to the trimeric assembly, see Figure 1a) and (ii) that all rungs are equivalents, which is not true since the rungs at the extremities have particular roles.^29,31^ Indeed, the rungs at the N-ter and C-ter of the helix have key roles in nucleating and terminating the fold. As such, they belong to the so-called “capping motifs” at the extremities of the helix that are the structural elements, belonging to the β–helix or not, responsible for the solubility of the protein.^7,33–36^ It has indeed been shown that removing capping motifs of β–solenoids, including left-handed β–helix, allows their self-assembly into fibers.^22,23,27^ While these aspects were considered by Balaram and coworkers first, and later by Prabha and Balaji, they did not provide a specific analysis of the sequence of these N-terminal and C-terminal rungs.^30,36^

In this present work, we propose a sequence analysis of LβH-I by taking into account simultaneously that (i) LβH-I have three different faces due to the trimeric arrangement and that (ii) LβH-I is composed of three rungs of different nature. It has been performed only with sequences extracted from proteins of which the crystal structure in available in the Protein Data Bank (PDB), hence leaving no doubt on the position occupied in the LβH-I fold by each considered amino-acid. This analysis also helps identifying and understanding the role of specific motifs in initiating, stabilizing or terminating the fold. Overall, this study aims at (i) improving our understanding of the LβH-I’s sequence-structure relationship, and at (ii) providing guidelines for the rational design of protein sequences able to adopt a stable LβH-I fold.

## Results

In order to better understand the sequence-structure relationship of LβH-containing proteins, we could identify 78 different crystallographic structures (complete list Table S1) querying UniProt IDs in the PDB. The analysis procedure that was followed is illustrated Figure 1. First, for each structure, all the parts of the protein not belonging to the LβH fold were omitted. As a consequence, all interactions between the canonical parts of LβH and other parts of the proteins have not been considered in this study. Second, the three different faces of the helix within the trimer were discriminated (Figure 1c). They were labelled according to their solvent exposition: buried (b), exposed (e) and partially exposed (p).^36^ They are composed of six residues, hence termed hexad, four for the β–strand (B1, B2, B3, B4), and 2 for the turn (T1, T2).^29^Of note, in positions B1 and B3, the residues are oriented inwards, hence forming the hydrophobic core, whereas the others are oriented outwards of the helix (Figure 1d).. Third, the three different parts of the helix, the N-terminal capping rung (N), the Inner rungs (I), and the C-terminal capping rung (C) were distinguished (Figure 1d). As a consequence, the resulting analysis reports 9 different hexads repeats, which can be presented in a table of 3 lines (parts of the helix) and 3 columns (sides of the helix) (Figure 1e). The full table is given Figure S1, but for clarity sake, in the following main text, it will be presented line per line.

The result of this analysis is the position-dependent residue propensity for each hexad. In other words, for the different positions, each amino acid is depicted as its letter, and the size of the latter is proportional to the occurrence of the amino acid (Fig. 2, 5 and 7).

Hereafter, each residue’s position will be referred to as its position within its hexad: eg B3(C.e) refers to the residue in position B3 of the hexad located in the C-terminal Capping rung (C) of the helix, in the face of the helix that is the most exposed to the solvent (e).

### Analysis of the inner rungs

The inner rungs are the most represented in the analysis, with more than 72 % of residues, as they constitute the core of the helix. We observe that the three hexads of inner rungs are relatively close in composition (Figure 2), explaining their proximity with results reported by Cohen and coworkers.^37^ Yet, our analysis reveals some differences, notably at the T1 and T2 positions.

**Figure 2.**
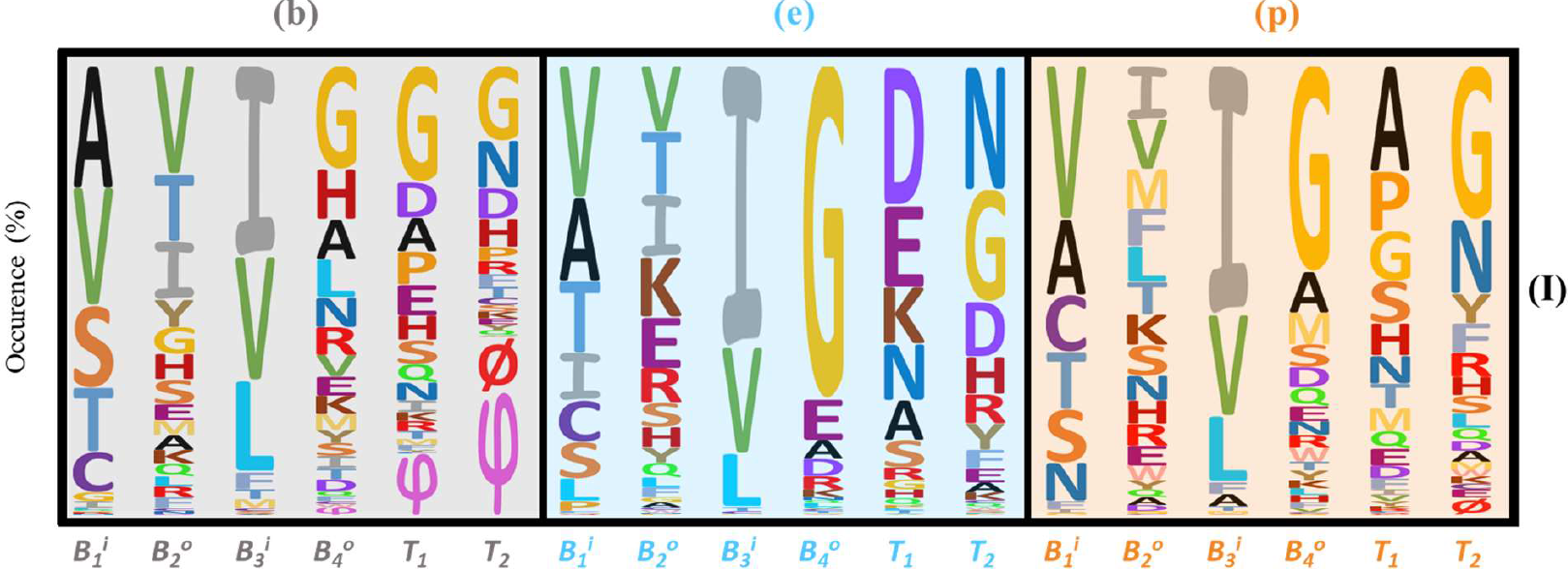
Occurrence of residues for each one of the eighteen positions of the inner rungs. The height of the residue’s letter is proportional to its frequency. φ : residues involved in a loop or a non-canonical conformation ; ∅ : missing residues.

The residues composing the hydrophobic core, at B_1_ and B_3_ positions, are very similar for the three faces (Figure 2). This is expected since, because they are buried, they should be the least impacted by the differences in solvent-exposition of the faces.

B3 position is occupied almost exclusively by Ile, Val and Leu (Figure 2). Even though Ile is the most frequent for all 3 faces, only 16 % of the inner rungs contain 3 Ile (at B3 (b), (e) and (p)). Instead, the most frequent configurations are (Ile)2(Val/Leu)1 and (Ile)1(Val)1(Leu)1 (Figure S3).

While Valine and Alanine are the most frequent at B1 position, residues with an alcohol or a thiol group (Ser, Thr, Cys) are also commonly encountered (Figure 2, S4 and S5). Indeed, these group make H-bonds with the main chain of residues i-3 (ie B4) and i+18 (ie same B1 position, in the following rung) (Figure 3a). Plus, in the case of Thr, the methyl group of its side chain also participates to the hydrophobic core. Interestingly, even if the residue B1 is usually part of the hydrophobic core, a non-negligible proportion of Asn is found at the B1(I.p), ca. 10%. Internal Asn ladders have been observed, also interacting with the main chain of B4 (e) (Figure 3b), and evidenced by tables analysing the interdependence of residue composition of adjacent rungs (Figure S8). This is reminiscent to the Asn ladders observed and described in the less regular β– helix of PelC and PelE.^38^ It is also interesting to observe Pro at B1(I.p) positions, as it is normally “forbidden” at this position.31 Actually, it is observed only right after one of the loop that that are often observed at this corner, that are very variable in length and structure. In such case, the side chain of Pro is oriented towards the interior of the helix, contributing to the hydrophobic core while imposing a torsion on the main chain, bringing it back to the canonical fold (Figure 3c). Noteworthy, the presence of a loop prior the B1(I.p) position seems not compatible with the presence of Ser or Thr at this position, probably because of their hydrophilicity. It is then preferred to accommodate a Val, an Ile or a Pro.

**Figure 3.**
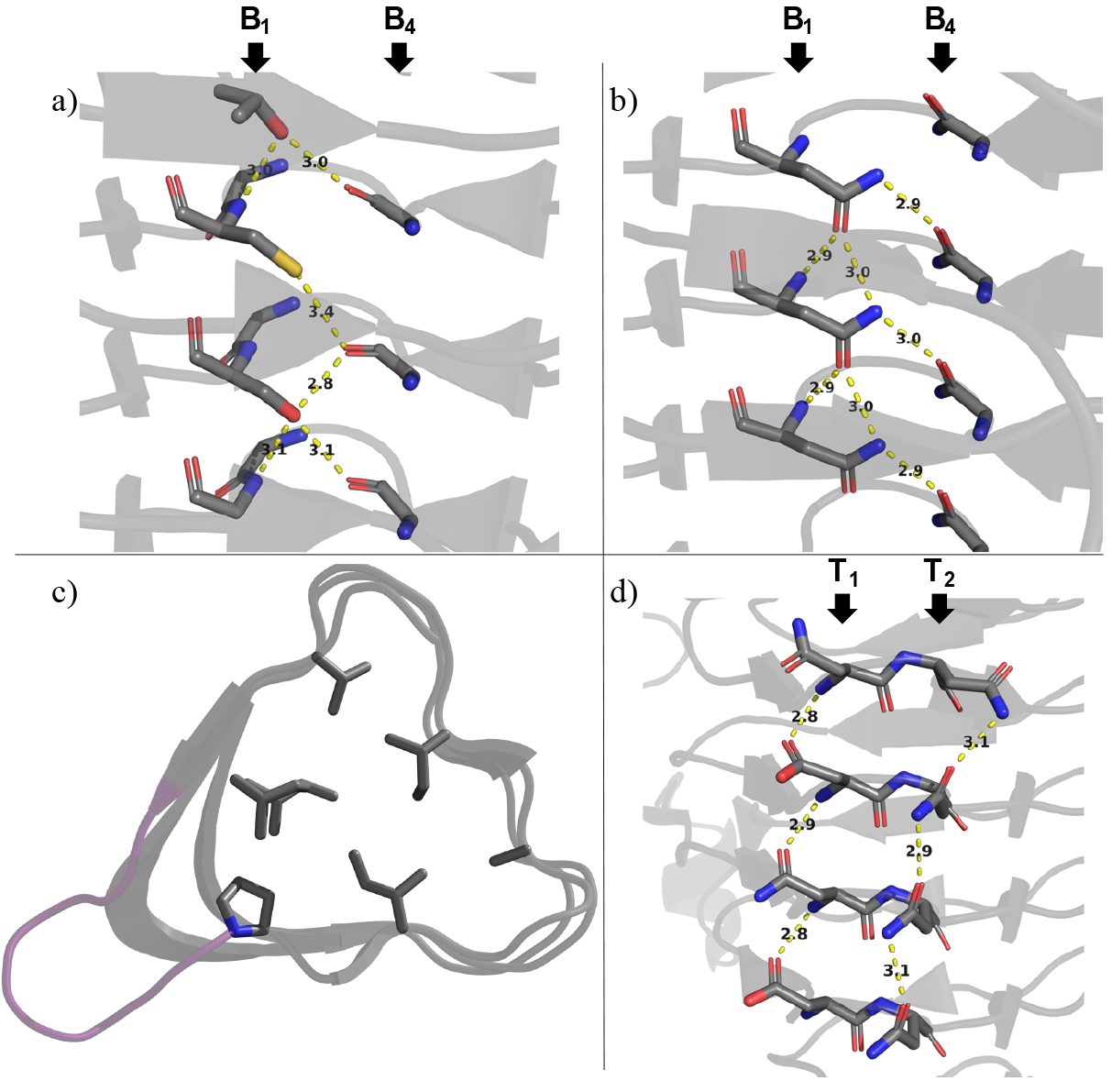
Illustration of stabilizing interactions observed in inner rungs. a) Thr, Ser and Cys in B1 can make H-bonding with the main chain of the amino acid, at the B4 position preceding it, the Thr methyl group oriented towards the hydrophobic core (PDB : 3IXC). b) An internal asparagine ladder (at the B1 position) interacting like the residues displayed in a) (PDB : 4EQY). c) Overview of the hydrophobic core (B1 and B3 side chains) and highlighting the Pro residue participating to the hydrophobic core after a loop (in pink) (PDB: 1XHD). d) Illustration of the hydrogen bonds involving Asp and Asn positions T1 and T2 of rung (I), face (e) (PDB: 1OCX).

In the face (e) that positions where side chain point outward the helix (ie B2, B4, T1 and T2) are mainly occupied by polar and charged residues while they are less frequent for the other two faces. Focusing on position B4, the main observations made previously are confirmed:^29,30^ (i) Glycine is frequent in this position, and (ii) especially at the face (e) (74% in this study). As can be seen from the Figure 2, B4(I.b) can also be the start of the variable loops observed at the corners (b-e), although these loops mainly substitute the positions T1-2(I.b) (Figure 2, φ symbols).

In the canonical rungs (I), prolines are scarce and found almost exclusively in positions T1. It accounts for 1%, 8% and 14% of the residues for the faces (e), (b) and (p), respectively. Interestingly, the latter (ie T1(I.p)) is at the interface of the helices in the trimers, together with T2(I.p) and B2(I.p), explaining why residues with short side chains are most often observed (Ala, Pro, Gly, Ser) (Figure 2).

Globally looking at the composition of the full rung (I), it appears clearly that polar residues are mainly located at positions T1(I.e) and T2(I.e), which corresponds to the corner well-exposed to the solvent, but which do not contain variable loops. Indeed, the most frequent residues there are Asp for T1(I.e) and Asn for T2(I.e). They both participate to the global stabilization of the fold as their side chain are involved in H-bonds with one or two adjacent rungs: at T1(I.e), Asp and Asn accept a H-bond of the amide (nitrogen) of the residue i-18, and at T2(I.e), Asn can form Asn ladders,^32^ thus interacting with residues i-18 and i+18 (Figure 3d).

The position T2(I.p) is located at the interface between the three helices, at the centre of the trimer. While Gly residues are often observed, probably to avoid clashes, Asn and aromatic residues are able there to perform stabilizing interactions, reinforcing the cohesion of the trimeric assembly (Figure 4). Interestingly, the alternating of Gly and Asn from one rung to the next, is the most frequent pattern observed at T_2_(I.p) (Figure S13).

**Figure 4.**
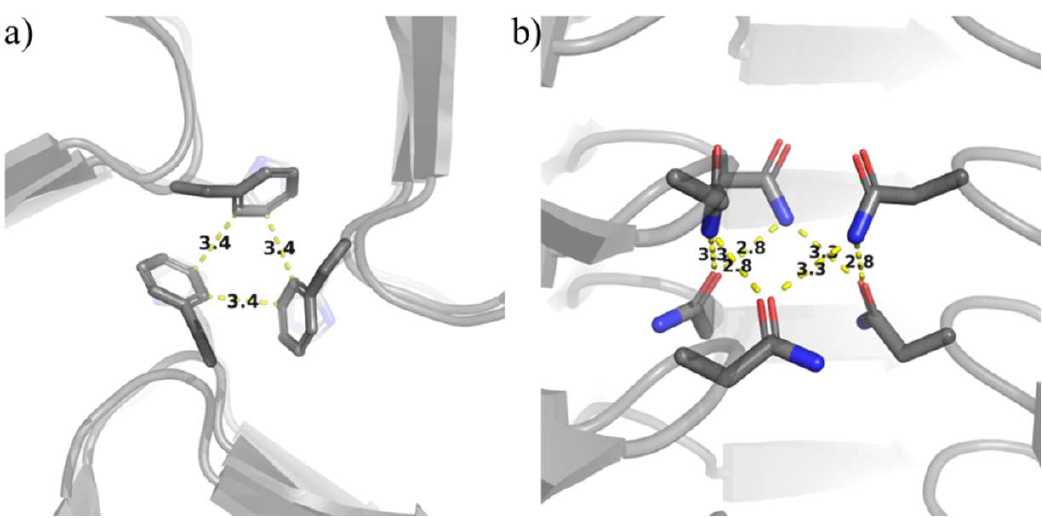
Examples of interactions located at the interface between the three helices, involving residues at position T2(I.p). a) Phe residues interactions (PDB: 1LXA), b) Asn ladders and interactions between helices (PDB: 4EQY)

Finally, at the B2 positions, the most frequent amino acids are β–branched, while one might have expected more polar residues as B2 is oriented outwards. The reason might be that β–branched residues are favourable for the formation of β–sheets.^39^ Of note, of the B2 positions of inner rungs, the face (e) is where long and polar residues (Lys, Glu, Arg) are most observed, together representing 32% of the residues (9 % for (b) and 16 % for (p)).

### Analysis of the rungs at the extremities of the LβH-I

In order to avoid aggregation of the β-helix, by intermolecular interactions via its ends, or even the uncontrolled continuation of the fold, the helix is capped at both N-ter and C-ter.34,35 Most of the times, the main capping element is a loop or non-repetitive structure, interacting with the end of the β-helix, covering and its main chain’s amides groups and its hydrophobic core, at least partially. On top of that, it has been observed that the first (N-ter) and last (C-ter) rungs of the β-helix also display specific structural motifs participating to its termination (ie prevent further β– interaction).34,35 Hereafter, they will be described into more details, and in particular, how they arise from the sequence. Of note, these first (N) and last (C) rungs, not only vary significantly compared to the rungs (I), but also from a β-helix to another.

### The N-terminal rung – promoting the nucleation of the β-helix

The N-terminal rung (N) already differs visually from the others, (I) and (C). Indeed, its turns are distorted, with a tendency to fold back towards the centre of the helix. Richardson & Richardson described them as upward protrusion, which help preventing continuation of β interactions towards N-terminus.^34^ In addition to this role, these turns might actually have a nucleation role, orienting the fold towards a LβH-I rather than more linear or less regular β–fold. In particular, we identified a specific triad, that both favours (i) the formation of a turn and (ii) the interaction with an adjacent rung (towards C-ter). The sequence of these nucleation motifs is X-Pro-Z (B_4_-T_1_-T_2_), where X = His, Asp or Glu, and Z = Ser, Thr, Asp, or Asn (Figures 5 and 6). While Pro induces a structural constrain onto the main chain, hence forcing the formation of a turn, the side chains of X and Z form a hydrogen bond together, thus narrowing the angle of the turn. In addition, the side of Z forms a hydrogen bond with the main chain amide of the residue (i+18), which favours the formation of the helix. These triads are mostly found at (b) and (p), resulting in Pro being the most frequent residue at T1(N.b) and T1(N.p).

**Figure 5.**
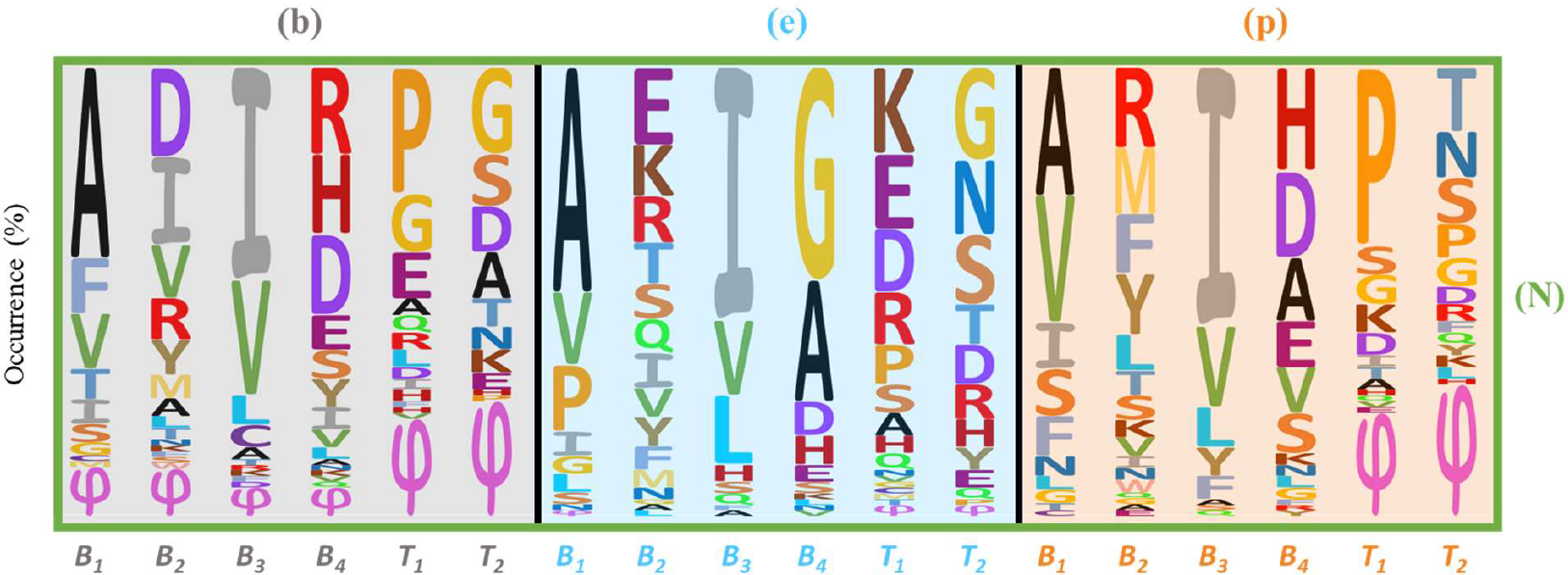
Occurrence of residues for each one of the eighteen positions in the N-terminal rung (N). The height of the residue’s letter is proportional to its frequency. φ : residues involved in a loop or a non-canonical conformation ; ∅ : missing residues.

**Figure 6.**
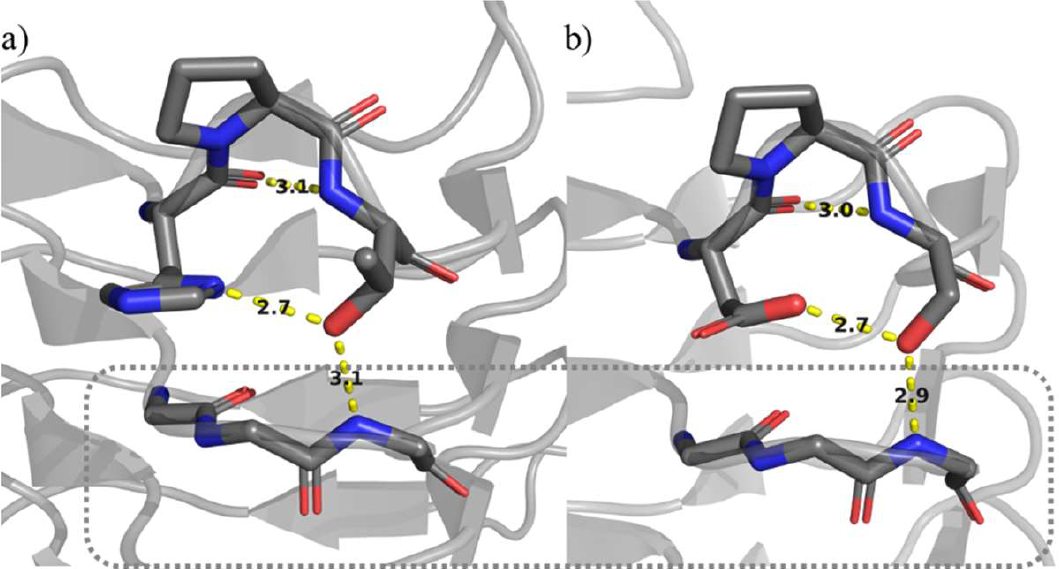
Illustrations of nucleation motif Xxx-Pro-Zzz in a rung (N). a) His-Pro-Thr (PDB: 3HSQ) and c) Asp-Pro-Ser (PDB: 4E6U). The dashed square highlights the rung n+1, ie the first inner rung (I).

Compared to the rungs (I), the residues compositing the hydrophobic core are similar, yet being more often composed of three Ile at B3 positions (Figure S3). Only in B1(N.e) one could note a higher proportion of Pro, which often occurs when the backbone returns to the canonical LβH-I fold, as shown before with the (I) rungs (*vide supra*).

On the other hand, the positions B2 differ from (I) to (N) as we observe here a higher proportion of non-β-branched or of polar residues. Another difference with rungs (I) is the presence of loops (visible as Phi symbols in Figure 5) at T1,2(N.p) in addition to T1,2(N.b).

Of note, there is no clear preference for a particular face for the start of the LβH-I fold. Indeed, for roughly half of the β–helices analysed here, the first (N) β–strand was of the face (p), and one fourth for faces (b) and (e).

### The C-terminal rung – ending the fold

Unlike the (N) rung, of which structural features might prevent aggregation on the N-ter of the helix while helping the proper LβH-I fold to initiate, the particular features observed for rung (C) only explain its termination role. When looking at the structures and their sequence, three main features are prominent.

First, the absence (empty set symbol in Figure 7) of residues at two positions, T_1-2_(C.b), resulting in a shorter loop for c.a. 70 % of the helices analysed here (Figure 8a). This diminishes the number of H-bond accessible on the C-terminal end, and also constrains the hydrophobic core to be smaller than in the rest of the helix, resulting in higher proportion of Val and Ala at positions B_3_(C). Indeed, observing three Val at the B3 positions of rung (C) is significantly more common than in other rungs (Figure S3).

**Figure 7.**
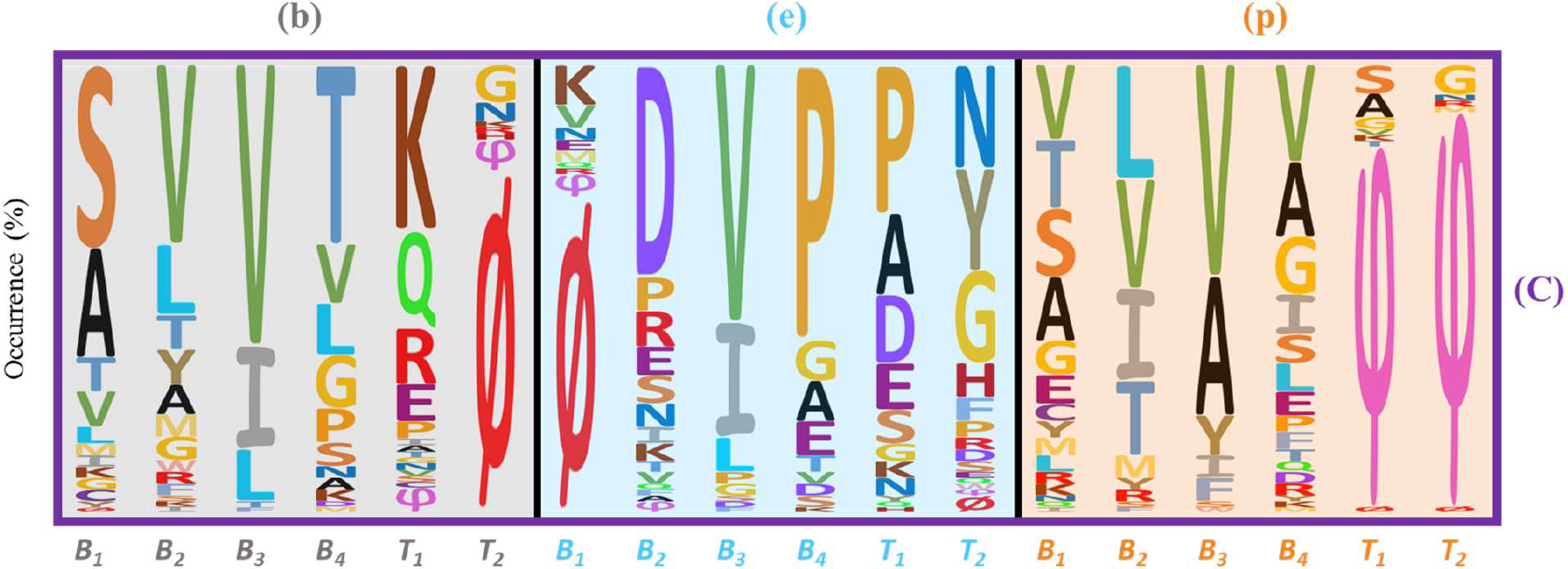
Occurrence of residues for each one of the eighteen positions in the C-terminal rung (N). The height of the residue’s letter is proportional to its frequency. φ : residues involved in a loop or a non-canonical conformation ; ∅ : missing residues.

**Figure 8.**
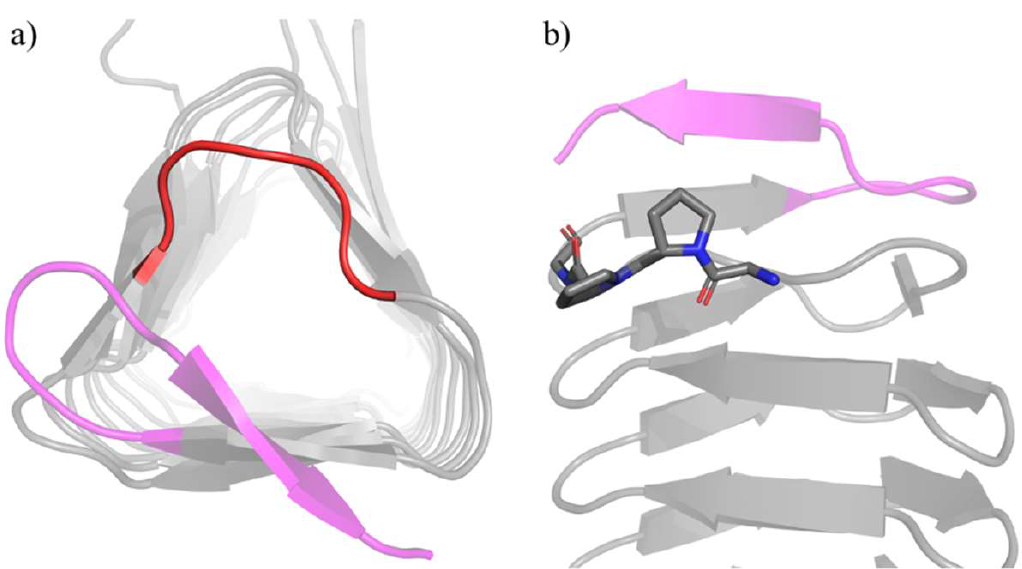
Characteristic motifs observed in rungs (C) illustrated at the C-terminal end of the UDP-N-acetylglucosamine acyltransferase (PDB: ILXA). a) Deletion of residues at positions T1,2(C.b) results in a shorter corner (in red). b) Presence of the Pro-Pro motif at the B4-T1(C.p), with the side-chain of the first oriented along the axis of the helix.

Second, the presence of a Pro in position B4(C.e) in c.a. 60 % of the structures analyzed, often accompanied with a Pro in the following T1 position (Figure 7). As evidenced in the past, a Pro in position B4 is not compatible with the LβH-I fold.^31^ It is then used in this C-terminal rung as a “helix breaker” to prevent the formation of a new β sheet by steric interaction, as the side chain of Pro at this position is oriented in the same axis as the helix (Figure 8b) hence sterically forbidding further β interaction.

Third, on the contrary to the start of the LβH-I, its termination has a marked preference for a face, with ca 90 % of the analyzed structures ending in face (p) (Table S2). In such case the last occupied position of the canonical fold is B4(C.p) (Table 7). The following residues are the start of the capping loop, which often (in ca 75 % of the observed structures, Table S2) was forming a β– hairpin with the final β–strand (C.p). Although being beyond the scope of this study as it does not belong to the LβH-I fold *per se*, the residue composition of this hairpin of the structures analyzed here is provided Figure S6. As a last remark, these C-ter capping motifs can further stabilize the trimer of β–helices, either by covering the hydrophobic core of another β–helix, or by further folding as an α–helix along the groove formed by two β–helices (Figure S40).

## Discussion

The analysis presented herein is not a classical bioinformatics study performed on a large set of LβH-I. Instead, it was limited to those of which the crystal structure was available in the PDB. As a consequence, this analysis suffers from biases inherent to the narrow scope and redundancy of the sequences studied (majority of them belong to the family of UDP N-Acetyltransferase). This doesn’t affect its relevance regarding the comprehension of the sequence-structure relationship, but limits its representativeness of the LβH-I-containing proteins. Its aim however was to provide a better understanding of the sequence-structure relationship for the LβH-I fold, based on structural features that are established with certainty by X-ray structure visualization. This allowed to identify interesting structural features, even if not present in a majority of sequences, such as the presence of Pro in B1(I.e) after a loop for example. In addition, we considered (i) the different nature of the different types of rung as well as (ii) the solvent exposure of the different face of the helix within the homo-trimer. This is the first time these two aspects are taken together into account during the analysis of LβH-I, thus distinguishing nine hexads. It first confirmed all previous observations, and second allowed the new description of specific motifs such as the Xxx-Pro-Zzz triad in the rung (N) for example, revealing the importance of this latter in initiating the LβH-I fold. While being clearly part of the β–helix, this work clearly shows that both rungs (N) and (C) should be considered as being part of the so-called “capping motif” as they possess structural features that are clearly involved in preventing the continuation of the LβH-I fold. This is in line with Richardson & Richardson viewpoint, but not with that of Bryan *et al*. as these rungs are ending or starting with an element of regular secondary structure.^34,35^

Complementary information, especially concerning the succession of residues from a rung to another at one particular position, were not discussed in the main text but are given the supplementary information (Figures S7 to S40).

Finally, to people aiming at designing LβH-I proteins, we advise to base their sequence design on the structural features highlighted above while following these general principles:

- In the most conserved positions, restrictions must be respected while optimizing the stability of the hydrophobic core and the number of H-bonds involving between adjacent rungs.
- Use the positions where a higher diversity of residues is allowed to introduce stabilizing interactions between adjacent rungs (eg Asn ladders or electrostatic interaction Lys/Glu), and, to introduce charged residues in order to provide the entire protein with a net charge to ensure its solubility at the desired pH range. In other words, adjust its pI according to the final application of the protein.
- Optimise the sequences of (N) and (C) rungs to initiate and terminate the LβH-I folding, respectively.

## Conclusion

This detailed analysis of the LβH-I fold is a significant step forward in our understanding of its sequence-structure relationship. Indeed, the analysis we provide reflects the trimeric nature of LβH-Is, as well as the nucleation and termination roles of their extremities, with a higher fidelity than before. It thus paves the way for the rational design of peptides and proteins folding into LβH-I, therefore particularly relevant for fields such as artificial protein design or bottom-up biomaterial synthesis.

## Materials and methods

All the protein crystal structures analysed here (Table S1) have been downloaded from the Protein Data Bank (PDB). The Uniprot code was used to search for structures and a similarity search has been performed to find similar other structures. The Database of Structural Repeats in Proteins (DbStRiP) developed by Chakrabarty and Parekh has also been used as a source of information .4 PyMOL software was used to visualize all 78 structures and to generate pictures.40 For each LβH-I, the labelling of the faces and the rung types as well as the position annotation for each amino-acid were performed manually, following the methodology illustrated in Figure 1. Excel 2016 (Microsoft) was used to process data regarding sequences.

## Acknowledgments

Authors thank the Agence Nationale de la Recherche for funding (ANR-18-CE07-0022-01). They also thank O. Sénèque, M. Paternostre, W. Ghattas, V. Forge and C. Ménard-Moyon for helpful discussions.

## References

(1) Anfinsen, C. B. Principles That Govern the Folding of Protein Chains. Science 1973, 181 (4096), 223–230. https://doi.org/10.1126/SCIENCE.181.4096.223.

(2) Kajava, A. V; Steven, A. C. B-Rolls, B-Helices, and Other B-Solenoid Proteins. In Advances in Protein Chemistry; Elsevier, 2006; Vol. 73, pp 55–96.

(3) Kajava, A. V. Tandem Repeats in Proteins: From Sequence to Structure. J. Struct. Biol. 2012, 179 (3), 279–288. https://doi.org/10.1016/J.JSB.2011.08.009.

(4) Chakrabarty, B.; Parekh, N. DbStRiPs: Database of Structural Repeats in Proteins. Protein Sci. 2021. https://doi.org/10.1002/pro.4052.

(5) Tripp, K. W.; Barrick, D. The Tolerance of a Modular Protein to Duplication and Deletion of Internal Repeats. J. Mol. Biol. 2004, 344 (1), 169–178. https://doi.org/10.1016/J.JMB.2004.09.038.

(6) Javadi, Y.; Itzhaki, L. S. Tandem-Repeat Proteins: Regularity plus Modularity Equals Design-Ability. Curr. Opin. Struct. Biol. 2013, 23 (4), 622–631. https://doi.org/10.1016/j.sbi.2013.06.011.

(7) Parmeggiani, F.; Huang, P. S. Designing Repeat Proteins: A Modular Approach to Protein Design. Curr. Opin. Struct. Biol. 2017, 45, 116–123. https://doi.org/10.1016/J.SBI.2017.02.001.

(8) Paladin, L.; Bevilacqua, M.; Errigo, S.; Piovesan, D.; Mičetić, I.; Necci, M.; Monzon, A. M.; Fabre, M. L.; Lopez, J. L.; Nilsson, J. F.; Rios, J.; Menna, P. L.; Cabrera, M.; Buitron, aM. G.; Kulik, M. G.; Fernandez-Alberti, S.; Fornasari, M. S.; Parisi, G.; Lagares, A.; Hirsh, L.; Andrade-Navarro, M. A.; Kajava, A. V.; Tosatto, S. C. E. RepeatsDB in 2021: Improved Data and Extended Classification for Protein Tandem Repeat Structures. Nucleic Acids Res. 2021, 49 (D1), D452–D457. https://doi.org/10.1093/nar/gkaa1097.

(9) Pellegrini, M. Tandem Repeats in Proteins: Prediction Algorithms and Biological Role. Front. Bioeng. Biotechnol. 2015, 3 (SEP), 155885. https://doi.org/10.3389/FBIOE.2015.00143/BIBTEX.

(10) Jorda, J.; Kajava, A. V. T-REKS: Identification of Tandem REpeats in Sequences with a K-MeanS Based Algorithm. Bioinformatics 2009, 25 (20), 2632–2638. https://doi.org/10.1093/bioinformatics/btp482.

(11) Voet, A. R. D.; Noguchi, H.; Addy, C.; Simoncini, D.; Terada, D.; Unzai, S.; Park, S. Y.; Zhang, K. Y. J.; Tame, J. R. H. Computational Design of a Self-Assembling Symmetrical β-Propeller Protein. Proc. Natl. Acad. Sci. U. S. A. 2014, 111 (42), 15102–15107. https://doi.org/10.1073/pnas.1412768111.

(12) Norn, C. H.; André, I. Computational Design of Protein Self-Assembly. Curr. Opin. Struct. Biol. 2016, 39, 39–45. https://doi.org/10.1016/J.SBI.2016.04.002.

(13) Correnti, C. E.; Hallinan, J. P.; Doyle, L. A.; Ruff, R. O.; Jaeger-Ruckstuhl, C. A.; Xu, Y.; Shen, B. W.; Qu, A.; Polkinghorn, C.; Friend, D. J.; Bandaranayake, A. D.; Riddell, S. R.; Kaiser, B. K.; Stoddard, B. L.; Bradley, P. Engineering and Functionalization of Large Circular Tandem Repeat Protein Nanoparticles. Nat. Struct. Mol. Biol. 2020 274 2020, 27 (4), 342–350. https://doi.org/10.1038/s41594-020-0397-5.

(14) Vanderstraeten, J.; Briers, Y. Synthetic Protein Scaffolds for the Colocalisation of Co-Acting Enzymes. Biotechnol. Adv. 2020, 44, 107627. https://doi.org/10.1016/J.BIOTECHADV.2020.107627.

(15) Leibly, D. J.; Arbing, M. A.; Pashkov, I.; Devore, N.; Waldo, G. S.; Terwilliger, T. C.; Yeates, T. O. A Suite of Engineered GFP Molecules for Oligomeric Scaffolding. Structure 2015, 23 (9), 1754–1768. https://doi.org/10.1016/j.str.2015.07.008.

(16) Luo, Q.; Hou, C.; Bai, Y.; Wang, R.; Liu, J. Protein Assembly: Versatile Approaches to Construct Highly Ordered Nanostructures. Chem. Rev. 2016, 116 (22), 13571–13632. https://doi.org/10.1021/acs.chemrev.6b00228.

(17) Gwyther, R. E. A.; Dafydd Jones, D.; Worthy, H. L. Better Together: Building Protein Oligomers Naturally and by Design. Biochem. Soc. Trans. 2019, 47 (6), 1773–1780. https://doi.org/10.1042/BST20190283.

(18) Zhu, J.; Avakyan, N.; Kakkis, A.; Hoffnagle, A. M.; Han, K.; Li, Y.; Zhang, Z.; Choi, T. S.; Na, Y.; Yu, C. J.; Tezcan, F. A. Protein Assembly by Design. Chem. Rev. 2021, 121 (22), 13701–13796. https://doi.org/10.1021/ACS.CHEMREV.1C00308.

(19) Plückthun, A. Designed Ankyrin Repeat Proteins (DARPins): Binding Proteins for Research, Diagnostics, and Therapy. Annu. Rev. Pharmacol. Toxicol. 2015, 55, 489–511. https://doi.org/10.1146/annurev-pharmtox-010611-134654.

(20) Urvoas, A.; Valerio-Lepiniec, M.; Minard, P. Artificial Proteins from Combinatorial Approaches. Trends Biotechnol. 2012, 30 (10), 512–520. https://doi.org/10.1016/j.tibtech.2012.06.001.

(21) Deng, X.; Gonzalez Llamazares, A.; Wagstaff, J. M.; Hale, V. L.; Cannone, G.; McLaughlin, S. H.; Kureisaite-Ciziene, D.; Löwe, J. The Structure of Bactofilin Filaments Reveals Their Mode of Membrane Binding and Lack of Polarity. Nat. Microbiol. 2019, 4 (12), 2357–2368. https://doi.org/10.1038/s41564-019-0544-0.

(22) Scotter, A. J.; Guo, M.; Tomczak, M. M.; Daley, M. E.; Campbell, R. L.; Oko, R. J.; Bateman, D. A.; Chakrabartty, A.; Sykes, B. D.; Davies, P. L. Metal Ion-Dependent, Reversible, Protein Filament Formation by Designed Beta-Roll Polypeptides. BMC Struct. Biol. 2007, 7 (1), 63. https://doi.org/10.1186/1472-6807-7-63.

(23) Peralta, M. D. R. R.; Karsai, A.; Ngo, A.; Sierra, C.; Fong, K. T.; Hayre, N. R.; Mirzaee, N.; Ravikumar, K. M.; Kluber, A. J.; Chen, X.; Liu, G.-Y. Y.; Toney, M. D.; Singh, R. R.; Cox, D. L.; Hayre, R.; Mirzaee, N.; Ravikumar, K. M.; Chen, X.; Liu, G.-Y. Y.; Toney, M. D.; Singh, R. R.; Lee, D.; Hayre, N. R.; Mirzaee, N.; Ravikumar, K. M.; Kluber, A. J.; Chen, X.; Liu, G.-Y. Y.; Toney, M. D.; Singh, R. R.; Cox, D. L. Engineering Amyloid Fibrils from β-Solenoid Proteins for Biomaterials Applications. ACS Nano 2015, 9 (1), 449–463. https://doi.org/10.1021/nn5056089.

(24) Altamura, L.; Horvath, C.; Rengaraj, S.; Rongier, A.; Elouarzaki, K.; Gondran, C.; Maçon, L. B.; Vendrely, C.; Bouchiat, V.; Fontecave, M.; Mariolle, D.; Rannou, P.; Le Goff, A.; Duraffourg, N.; Holzinger, M.; Forge, V. A Synthetic Redox Biofilm Made from Metalloprotein-Prion Domain Chimera Nanowires. Nat. Chem. 2017, 9 (2), 157–163. https://doi.org/10.1038/NCHEM.2616.

(25) Cheng, P.-N.; Pham, J. D.; Nowick, J. S. The Supramolecular Chemistry of β-Sheets. J. Am. Chem. Soc. 2013, 135 (15), 5477–5492. https://doi.org/10.1021/ja3088407.

(26) Baarda, R. A.; Marianchuk, T. L.; Toney, M. D.; Cox, D. L. In Silico Stress-Strain Measurements on Self-Assembled Protein Lattices. Soft Matter 2018, 14 (40), 8095–8104. https://doi.org/10.1039/C8SM00412A.

(27) Peng, Z.; Peralta, M. D. R.; Cox, D. L.; Toney, M. D. Bottom-up Synthesis of Protein-Based Nanomaterials from Engineered β-Solenoid Proteins. PLoS One 2020, 15 (2), e0229319. https://doi.org/10.1371/journal.pone.0229319.

(28) Yoder, M. D.; Keen, N. T.; Jurnak, F. New Domain Motif: The Structure of Pectate Lyase C, a Secreted Plant Virulence Factor. Science 1993, 260 (5113), 1503–1507. https://doi.org/10.1126/SCIENCE.8502994.

(29) Choi, J. H.; Govaerts, C.; May, B. C. H.; Cohen, F. E. Analysis of the Sequence and Structural Features of the Left-Handed β-Helical Fold. Proteins: Struct., Funct., Bioinf. 2008, 73 (1), 150–160. https://doi.org/10.1002/prot.22051.

(30) Iengar, P.; Joshi, N. V.; Balaram, P. Conformational and Sequence Signatures in β Helix Proteins. Structure 2006, 14 (3), 529–542. https://doi.org/10.1016/J.STR.2005.11.021.

(31) Choi, J. H.; May, B. C. H. H.; Govaerts, C.; Cohen, F. E. Site-Directed Mutagenesis Demonstrates the Plasticity of the β Helix: Implications for the Structure of the Misfolded Prion Protein. Structure 2009, 17 (7), 1014–1023. https://doi.org/10.1016/j.str.2009.05.013.

(32) Haspel, N.; Zanuy, D.; Alemán, C.; Wolfson, H.; Nussinov, R. De Novo Tubular Nanostructure Design Based on Self-Assembly of β-Helical Protein Motifs. Structure 2006, 14 (7), 1137–1148. https://doi.org/10.1016/j.str.2006.05.016.

(33) Courtemanche, N.; Barrick, D. The Leucine-Rich Repeat Domain of Internalin B Folds along a Polarized N-Terminal Pathway. Structure 2008, 16 (5), 705–714. https://doi.org/10.1016/J.STR.2008.02.015.

(34) Richardson, J. S.; Richardson, D. C. Natural β-Sheet Proteins Use Negative Design to Avoid Edge-to-Edge Aggregation. Proc. Natl. Acad. Sci. U. S. A. 2002, 99 (5), 2754–2759. https://doi.org/10.1073/PNAS.052706099/ASSET/642B4D64-016B-4107-8E47-2DD53796EC6A/ASSETS/GRAPHIC/PQ0527060006.JPEG.

(35) Bryan, A. W.; Starner-Kreinbrink, J. L.; Hosur, R.; Clark, P. L.; Berger, B. Structure-Based Prediction Reveals Capping Motifs That Inhibit β-Helix Aggregation. Proc. Natl. Acad. Sci. U. S. A. 2011, 108 (27), 11099–11104. https://doi.org/10.1073/PNAS.1017504108/SUPPL_FILE/PNAS.1017504108_SI.PDF.

(36) Prabha, A.; Balaji, P. V. Characterization of Left-Handed Beta Helix-Domains, and Identification and Functional Annotation of Proteins Containing Such Domains. Proteins: Struct., Funct., Bioinf. 2020, No. July, 1–15. https://doi.org/10.1002/prot.25990.

(37) Choi, J. H.; Govaerts, C.; May, B. C. H. H.; Cohen, F. E. Analysis of the Sequence and Structural Features of the Left-Handed β-Helical Fold. Proteins: Struct., Funct., Bioinf. 2008, 73 (1), 150–160. https://doi.org/10.1002/prot.22051.

(38) Yoder, M. D.; Lietzke, S. E.; Jurnak, F. Unusual Structural Features in the Parallel β-Helix in Pectate Lyases. Structure 1993, 1 (4), 241–251. https://doi.org/10.1016/0969-2126(93)90013-7.

(39) Pal, D.; Chakrabarti, P. β-Sheet Propensity and Its Correlation Withparameters Based on Conformation. Acta Crystallogr., Sect. D: Biol. Crystallogr. 2000, 56 (5), 589–594. https://doi.org/10.1107/S090744490000367X.

(40) Schrödinger, L.; DeLano, W. PyMOL. 2020.

