## Supplementary information for "A closer look at Type I left-handed β-helices provides a better understanding in their sequence-structure relationship: towards their rational design"

June 27, 2023

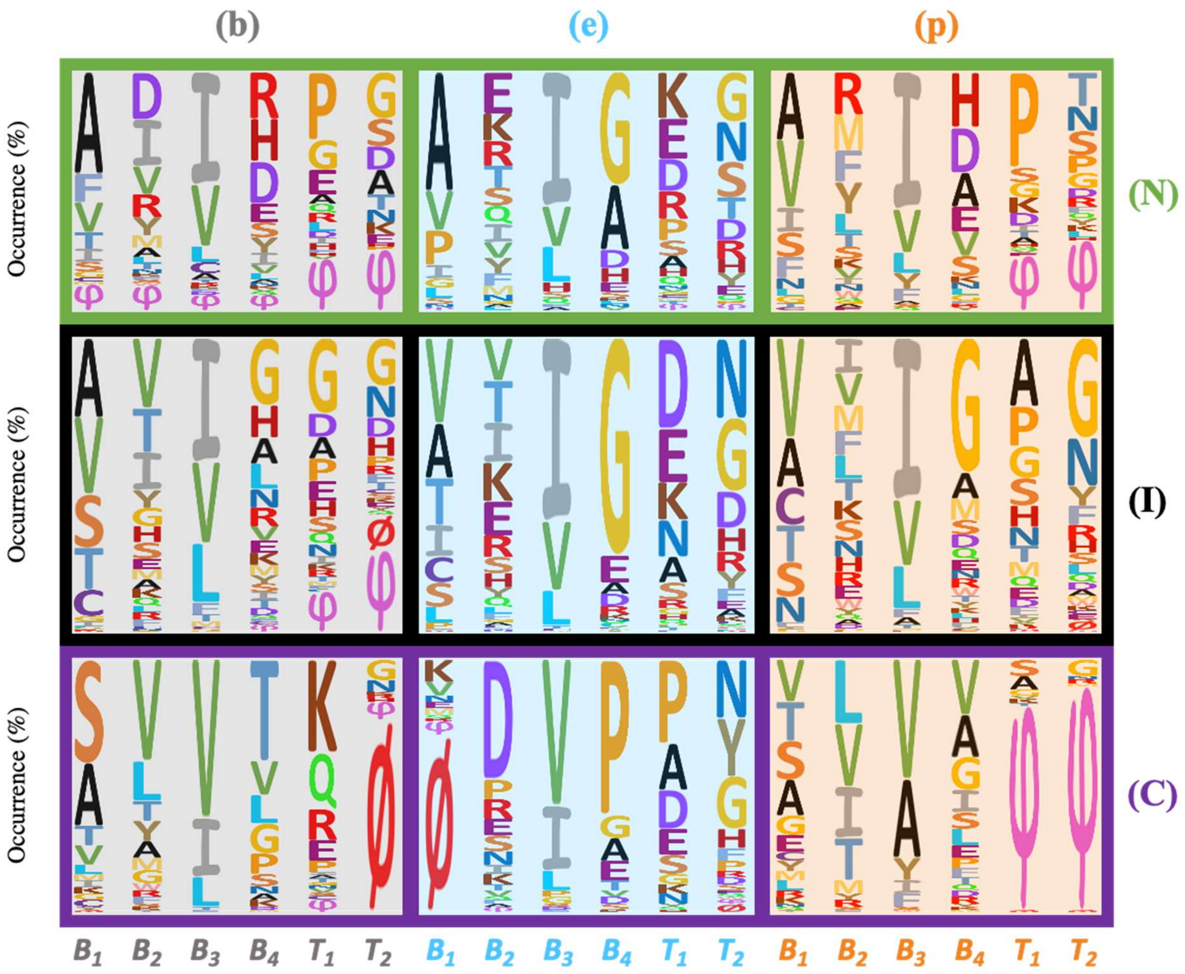


**Figure S1.** Occurrence of residues for each one of the 54 positions of the 9 hexads. The three columns are for the three faces (b, e and p) of the LβH-I in a homo-trimer. The three rows refer to the three types rungs (N, I, C) of the LβH-I. The height of the residue's letter is proportional to its frequency. The φ letter is referring to residues involved in a loop or a non-canonical conformation and Ø stands for missing residues.

**Table S1.** List of crystal structures analyzed in this study.

| PDB code (containing metal $) | Topology (mainly beta) CATH | Family type | Species (organism) | Number of rungs | Resolution (Ä) |
|---|---|---|---|---|---|
| **3OTM$** | γ-carbonic anhydrase | LYASE | Methanosarcina thermophila | 7 | 1.5 |
| **1LXA** | LpxA | ACYLTRANSFERASE | Escherichia coli K-12 | 10 | 2.6 |
| **4E6U** | LpxA | TRANSFERASE | Acinetobacter baumannii | 9 | 1.41 |
| **3PMO** | LpxD | TRANSFERASE | Pseudomonas aeruginosa | 10 | 1.3 |
| **3KWE$** | γ-carbonic anhydrase | LYASE | Thermosynechococcus elongatus BP-1 | 7 | 1.1 |
| **1SSM** | SAT | TRANSFERASE | Haemophilus influenzae | 5 | 2.15 |
| **3MQH** | 3-N-acetyl transferase WlbB | TRANSFERASE | Bordetella petrii DSM 12804 | 7 | 1.43 |
| **4E79** | LpxD | TRANSFERASE | Acinetobacter baumannii | 10 | 2.66 |
| **4M9C** | WeeI acetyltransferase | TRANSFERASE | Acinetobacter baumannii | 6 | 2.1 |
| **3R3R$** | γ-carbonic anhydrase | TRANSFERASE | Salmonella enterica subsp. Enterica serovar Typhimurium | 7 | 1.2 |
| **3VNP$** | GK2848 | STRUCTURAL GENOMICS, gamma-carbonic anhydrase-like | Geobacillus kaustophilus HTA426 | 7 | 2.4 |
| **1HM0** | GLMU | TRANSFERASE | Streptococcus pneumoniae | 10 | 2.3 |
| **3FS8** | QdtC | TRANSFERASE | Thermoanaerobacterium thermosaccharolyticum | 11 | 1.7 |
| **3R8Y** | DapD | TRANSFERASE | Bacillus anthracis | 6 | 1.7 |
| **3Q1X** | SAT | TRANSFERASE | Entamoeba histolytica | 5 | 1.59 |
| **3F1X** | SAT | TRANSFERASE | Bacteroides vulgatus ATCC 8482 | 5 | 2 |
| **2GGO** | glucose-1-phosphate thymidylyltransferase | TRANSFERASE | Sulfurisphaera tokodaii | 8 | 1.8 |
| **1V3W$** | γ-carbonic anhydrase | TRANSFERASE | Pyrococcus horikoshii OT3 | 7 | 1.5 |
| **4N6A** | SAT | TRANSFERASE | Glycine max | 5 | 1.75 |
| **5U2K** | Galactoside O-acetyltransferase | TRANSFERASE | Staphylococcus aureus subsp. Aureus COL | 5 | 1.38 |
| **2RIJ** | DapD | TRANSFERASE | Campylobacter jejuni | 6 | 1.9 |
| **3R5D** | DapD | TRANSFERASE | Pseudomonas aeruginosa | 6 | 1.8 |
| **4EA9** | PerB | TRANSFERASE | Caulobacter vibrioides | 6 | 0.9 |
| **1FWY** | GlmU | TRANSFERASE | Escherichia coli | 4 | 2.3 |
| **5B04** | eIF2B | TRANSLATION | Schizosaccharomyces pombe 972h- | 6 | 2.99 |
| **4MZU** | FdtD | ISOMERASE, TRANSFERASE | Shewanella denitrificans OS217 | 7 | 2.2 |
| **4N27$** | γ-carbonic anhydrase | TRANSFERASE | Brucella abortus 2308 | 7 | 2.73 |
| **3BSY** | PglD | TRANSFERASE | Campylobacter jejuni | 6 | 1.8 |
| **1XHD** | Putative acetyltransferase | TRANSFERASE | Bacillus cereus ATCC 14579 | 7 | 1.9 |
| **1OCX** | MAT | TRANSFERASE | Escherichia coli | 6 | 2.15 |
| **1T3D** | SAT | TRANSFERASE | Escherichia coli | 5 | 2.2 |
| **2XAT** | PaXAT | ACETYLTRANSFERASE | Pseudomonas aeruginosa | 5 | 3.2 |
| **2P2O** | Maltose transacetylase | TRANSFERASE | Geobacillus kaustophilus HTA426 | 6 | 1.74 |
| **2V0J** | Glmu | TRANSFERASE | Haemophilus influenzae | 10 | 2 |
| **3DHO** | Streptogramin Acetyltransferase | TRANSFERASE | Enterococcus faecium | 5 | 1.8 |
| **3EEV** | VCA0300 | TRANSFERASE | Brucella abortus 2313 | 5 | 2.61 |
| **3EG4** | DapD | TRANSFERASE | Brucella suis | 6 | 1.87 |
| **3GOS** | DapD | TRANSFERASE | Yersinia pestis | 6 | 1.8 |
| **3GVD** | CysE | TRANSFERASE | Yersinia pestis | 5 | 2.4 |
| **3IGJ** | MAT | TRANSFERASE | Bacillus anthracis | 6 | 2.6 |
| **3MC4** | SAT | TRANSFERASE | Brucella abortus 2308 | 5 | 1.95 |
| **3SRT** | MAT | TRANSFERASE | Clostridioides difficile 630 | 6 | 2.5 |

| | | | | | |
|---|---|---|---|---|---|
| **3TIO$** | γ-carbonic anhydrase | TRANSFERASE | Escherichia coli K-12 | 7 | 1.41 |
| **1J2Z** | LpxA | TRANSFERASE | Helicobacter pylori | 9 | 2.1 |
| **6SC4$** | γ-carbonic anhydrase | METAL BINDING PROTEIN | 6sc7 candidate division MSBL1 archaeon SCGC-AAA259I09 | 7 | 2.6 |
| **1G97$** | GlmU | TRANSFERASE | Streptococcus pneumoniae | 10 | 1.96 |
| **3IXC$** | hexapeptide transferase | TRANSFERASE | Anaplasma phagocytophilum str. HZ | 7 | 1.61 |
| **4MFG$** | Putative Carbonic Anhydrase | TRANSFERASE | Clostridioides difficile 630 | 7 | 2 |
| **6JVU** | CysE | TRANSFERASE | Klebsiella pneumoniae MGH 78578 | 5 | 2.8 |
| **6OSS** | LpxA | TRANSFERASE | Proteus mirabilis HI4320 | 9 | 2.19 |
| **6GE9$** | GlmU | TRANSFERASE | Mycobacterium tuberculosis H37Ra | 10 | 2.26 |
| **3T57** | LpxA | TRANSFERASE | Arabidopsis thaliana | 10 | 2.1 |
| **4IHF** | LpxD | TRANSFERASE/LIPID BINDING PROTEIN | Escherichia coli K-12, Escherichia coli str. 'clone D i14 | 10 | 2.1 |
| **6IVE$** | γ-carbonic anhydrase | METAL BINDING PROTEIN | Thermus thermophilus HB8 | 7 | 2.3 |
| **5VMK** | GlmU | TRANSFERASE | Acinetobacter baumannii | 10 | 2.55 |
| **5JXX** | LpxA | TRANSFERASE | Moraxella catarrhalis BBH18 | 9 | 3 |
| **1KRR** | GAT | TRANSFERASE | Escherichia coli | 6 | 2.5 |
| **3TDT** | THDP | ACYLTRANSFERASE | Mycobacterium tuberculosis variant bovis | 6 | 2 |
| **5DEP** | LpxA | TRANSFERASE | Pseudomonas aeruginosa PA7 | 9 | 2.16 |
| **2IU8** | LpxD | TRANSFERASE | Chlamydia trachomatis | 10 | 2.2 |
| **6AMZ** | DapD | TRANSFERASE | Acinetobacter baumannii | 6 | 2.5 |
| **3BXY** | DapD | TRANSFERASE | Escherichia coli O157:H7 | 6 | 2 |
| **4FCE** | GlmU | TRANSFERASE | Yersinia pestis CO92 | 10 | 1.96 |
| **3VBP** | D94N | TRANSFERASE | Bacillus cereus SJ1 | 7 | 2.3 |
| **3HSQ** | LpxA | TRANSFERASE | Leptospira interrogans | 9 | 2.1 |
| **4M98** | PglB | TRANSFERASE | Neisseria gonorrhoeae FA 1090 | 6 | 1.67 |
| **3R0S** | LpxA | TRANSFERASE | Campylobacter jejuni subsp. jejuni NCTC 11168 = ATCC 700819 | 9 | 2.3 |
| **5F42** | LpxA | TRANSFERASE | Francisella tularensis subsp. novicida U112 | 9 | 2.06 |
| **5E3R** | DapD | TRANSFERASE | Corynebacterium glutamicum ATCC 13032 | 6 | 1.85 |
| **3FSX** | DapD | TRANSFERASE | Mycobacterium tuberculosis H37Rv | 6 | 2.15 |
| **4H7O** | SAT | TRANSFERASE | Vibrio cholerae O1 biovar El Tor str. N16961 | 5 | 2.17 |
| **6CKT** | DapD | TRANSFERASE | Legionella pneumophila subsp. pneumophila str. Philadelphia 1 | 6 | 1.8 |
| **3TK8** | DapD | TRANSFERASE | 3TK11Burkholderia pseudomallei 1710b | 6 | 1.8 |
| **4EQY** | LpxA | TRANSFERASE | Burkholderia thailandensis E264 | 9 | 1.8 |
| **3CJ8** | DapD | TRANSFERASE | Enterococcus faecalis | 6 | 1.95 |
| **4R36** | LpxA | TRANSFERASE | Bacteroides fragilis NCTC 9343 | 9 | 1.9 |
| **6MFK** | CAT | TRANSFERASE | lizabethkingia anophelis | 5 | 1.65 |
| **5UX9** | CAT | TRANSFERASE | Aliivibrio fischeri ES114 | 5 | 2.7 |

**Table S2.** General statistics on the β-helices analysed in this study.

| | | Buried (b) | Exposed (e) | Partially-exposed (p) | Total |
|---|---|---|---|---|---|
| Number of residues analyzed (incl. φ and Ø) | $N_{ter}$ (N) | **461** | **463** | **453** | **1377** |
| | Inner (I) | **2310** | **2388** | **2619** | **7317** |
| | $C_{ter}$ (C) | **455** | **468** | **465** | **1388** |
| Total residues (incl. φ and Ø) | | **3226** (32%) | **3319** (32.9%) | **3537** (35.1%) | **10082** |
| % Structures containing metals | | Total: **16.7 %** (13 out of 78); **Zn** (8); **Mg** (4); **Ni** (1) | | | |
| Number of rungs (N+I+C) per helix | | Max. = 11 ; Min. = 4 ; Average = 7 | | | **551** |
| By which hexad does the helix start ? | | 25.6 % (20) | 24.4% (19) | 50.0 % (39) | |
| By which hexad does the helix end ? | | 6.4% (5) | 1.3% (1) | 92.3% (72) | |
| Capping motif after rung $C_{ter}$ | β hairpin | 76.9 % (60) of β hairpin in semi-exposed | | | |
| | Other | 23.1 % (18) of chains and α helix | | | |

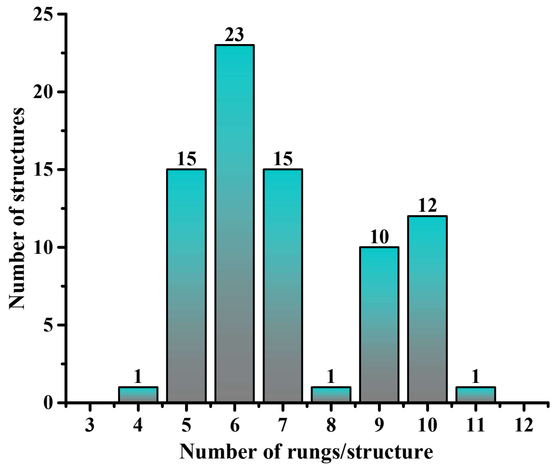


**Figure S2.** Diagram showing the number of rungs composing the different Type I left-handed β–helices considered in this study.

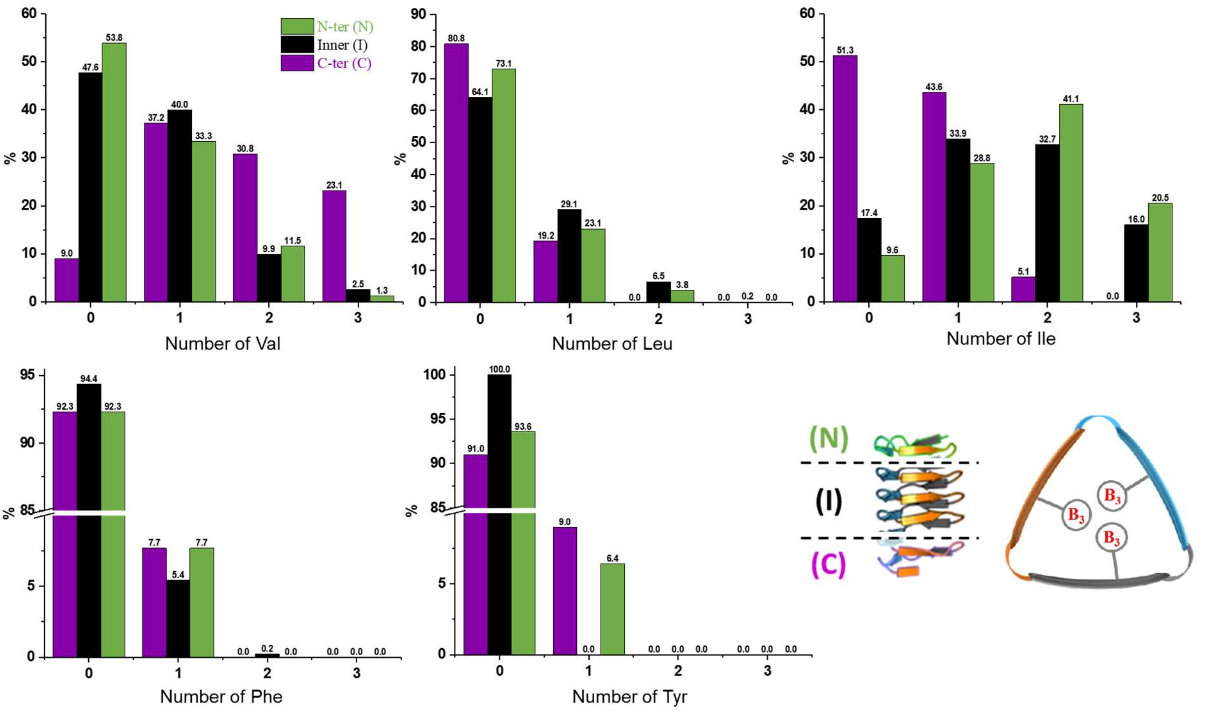


**Figure S3.** For a given amino acid, how many times does it appear in the $B_3$ positions of on particular rung. The schemes in the right bottom reminds the different types of rung, and where the $B_3$ positions are located in one rung.

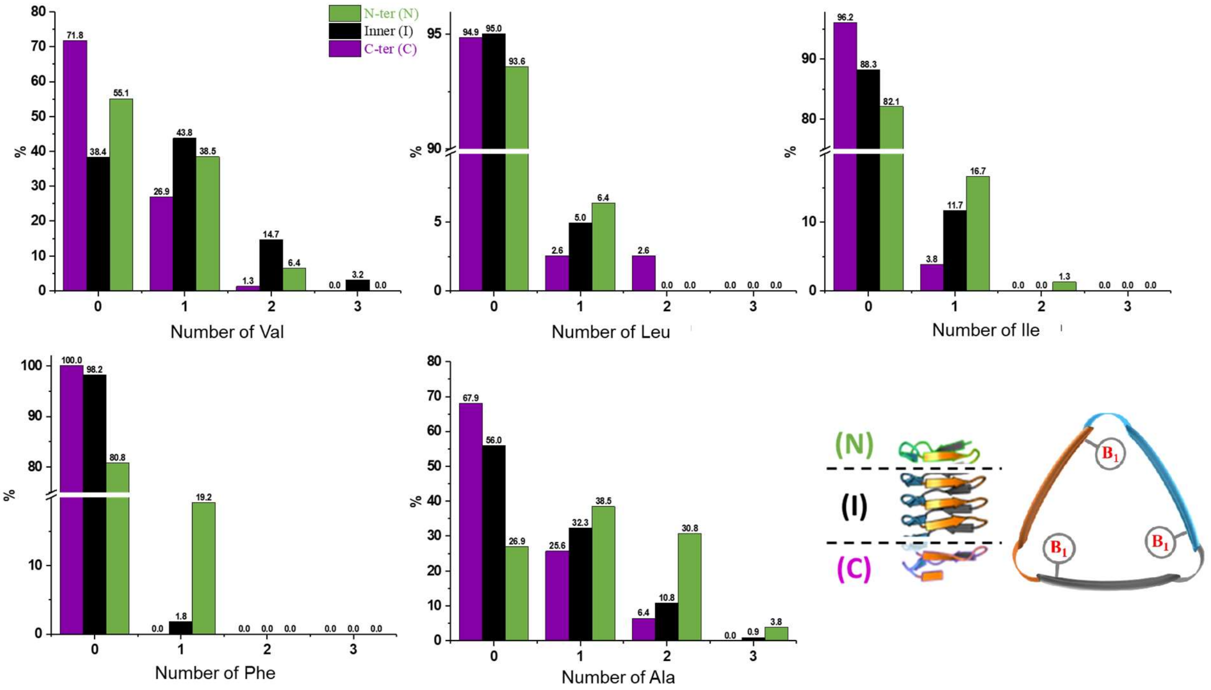


**Figure S4.** For a given amino acid, how many times does it appear in the $B_1$ positions of on particular rung. The schemes in the right bottom reminds the different types of rung, and where the $B_3$ positions are located in one rung.

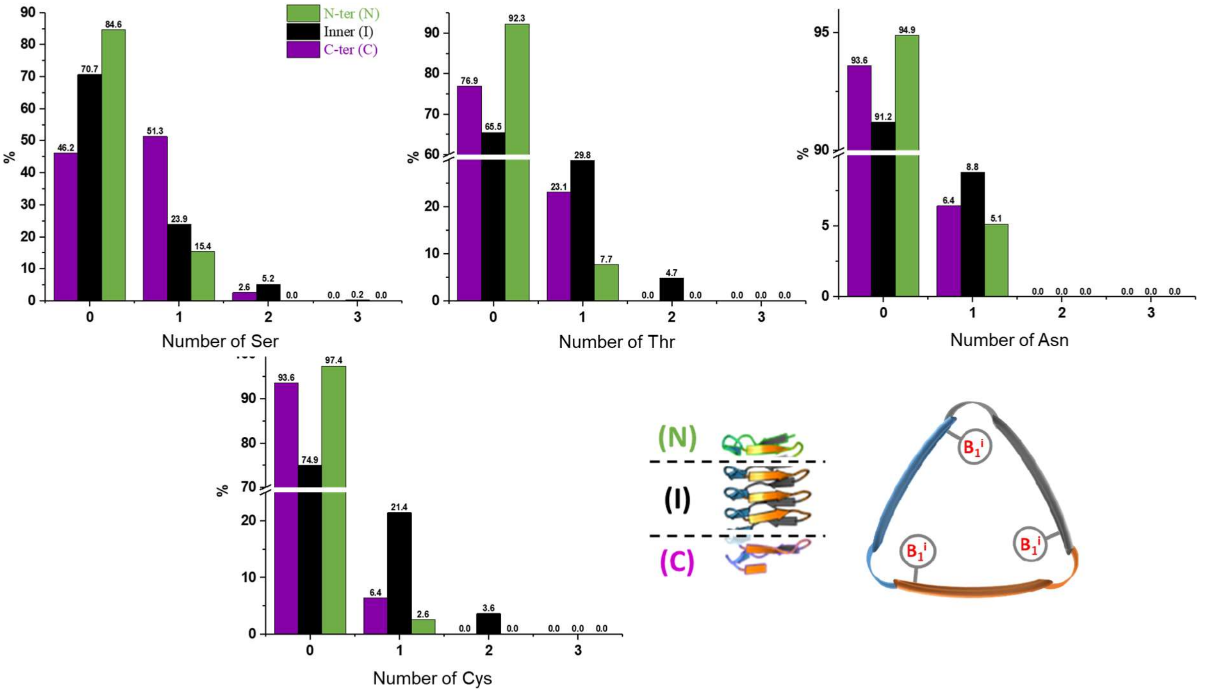


**Figure S5.** For a given amino acid, how many times does it appear in the $B_1$ positions of on particular rung. The schemes in the right bottom reminds the different types of rung, and where the $B_3$ positions are located in one rung.

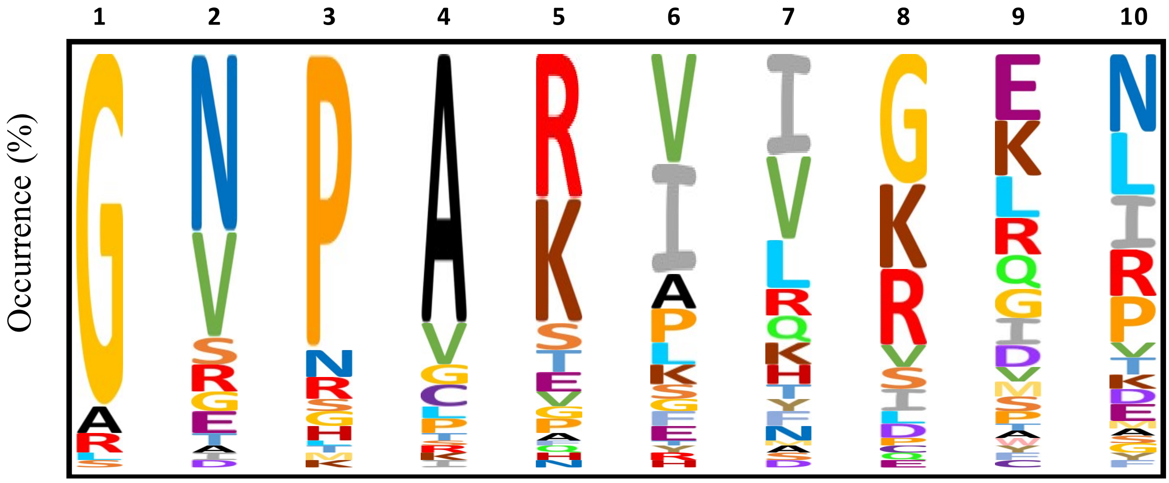


**Figure S6.** Occurrence of the residues in the second–strand of the capping β–hairpin. This sequence directly follows the end of the Type I left-handed β–helix, ie the position $B_4$(C.p).

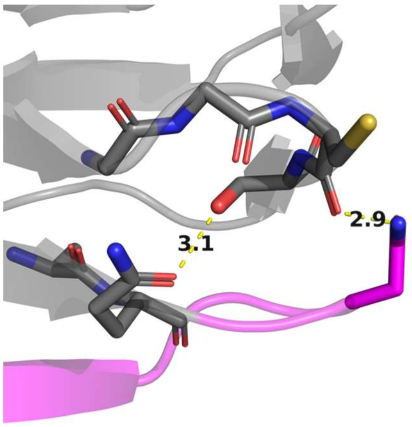


**Figure S7.** Stabilizing interaction observed at the C-terminal end of the helix, illustrated here with the UDP-N-acetylglucosamine acyltransferase (PDB: ILXA). Ser at $B_1$(I.b) position interacts with a Gln residue in $B_4$(C.p) and the Asn in the position 2 of the capping β–hairpin (in pink).

The following Figures (from S8 to S13) were generated the following way, using the Excel table, in which all the protein sequences have been entered into an 18-column table, with one line per rung. In order to analyze the **rung-to-rung succession of residues at one position within the same helix**, the Power Query has been used, and was made only for the Inner rungs. In a first column, has been copied the column corresponding to each position and defined as the source column. Then, this column has been indexed starting at 0 with increments of 1 through the formula “Table.AddIndexColumn”. The same operation has been done by indexing the previous column starting by 1 thanks to the same function. Then, the “Table.NestedJoin” has been used to join the rows of the 2 previous columns depending on the equality of values. The function “Table.ExpandTableColumn” has then been used to fuse the result of the previous operation with the source column that for the first and second indexed column. The “Table.UnpivotOtherColumns” function has been used to translate all the values into pairs of data referring to residues at rungs n and n+1. Finally, the final matrix has been generated through the “Table.pivot” function linked to a “List.Count” function to count the sequences of following amino acids. For each studied position, the resulting tables obtained for the 3 faces were pooled together in one Figure.

They can be read as the following example: “Figure S8 shows that, a Val has been observed 48 times at position $B_1$(I.b) (of the rung n) that was followed or preceded by an Ala at the same $B_1$(I.b) position in the rung n+1. We observed 50 times a Val at position $B_1$(I.p) that was followed/preceded by another Val at the same $B_1$(I.b) position in the following rung.”

b

| A.A | A | C | G | I | L | M | N | Q | R | S | T | V |
|---|---|---|---|---|---|---|---|---|---|---|---|---|
| A | 32 | | | | | | | | | | | |
| C | 8 | 3 | | | | | | | | | | |
| G | 2 | | 2 | | | | | | | | | |
| I | | | | | | | | | | | | |
| L | | | | | | | | | | | | |
| M | | 1 | | | | | | | | | | |
| N | | | | | | | | | | | | |
| Q | | | | | | | | | | | | |
| R | 1 | | | | | | | | | | | |
| S | 29 | 7 | 1 | | | | 1 | 2 | 1 | 5 | | |
| T | 13 | 9 | 1 | 1 | | | | | | 14 | 15 | |
| V | 48 | 22 | 1 | 7 | 2 | 1 | 1 | | | 33 | 27 | 16 |

e

| A.A | A | C | F | I | L | M | P | S | T | V |
|---|---|---|---|---|---|---|---|---|---|---|
| A | 21 | | | | | | | | | |
| C | 10 | | | | | | | | | |
| F | | | | | | | | | | |
| I | 3 | 4 | | 3 | | | | | | |
| L | 2 | 3 | | 2 | | | | | | |
| M | | | | | 1 | | | | | |
| P | 7 | | | 8 | | | | | | |
| S | 10 | 4 | | 1 | | | 2 | 8 | | |
| T | 17 | 18 | | 7 | 19 | | | 9 | 5 | |
| V | 18 | 21 | 2 | 27 | 5 | | 1 | 16 | 43 | 21 |

p

| A.A | A | C | F | G | I | N | S | T | V |
|---|---|---|---|---|---|---|---|---|---|
| A | 12 | | | | | | | | |
| C | 8 | 7 | | | | | | | |
| F | 1 | 3 | | | | | | | |
| G | 1 | | | | | | | | |
| I | 2 | | | | | | | | |
| N | 1 | 6 | | | 1 | 25 | | | |
| S | 11 | 19 | 1 | 1 | | 3 | 4 | | |
| T | 14 | 14 | | 2 | 1 | 11 | 8 | 4 | |
| V | 40 | 32 | 4 | 2 | 2 | 3 | 28 | 38 | 50 |

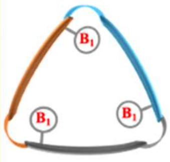


**Figure S8.** Rung-to-rung succession of residues at the $B_1$ position, for each face of the helix. The higher the number (indicated by intense red), the more frequent the succession.

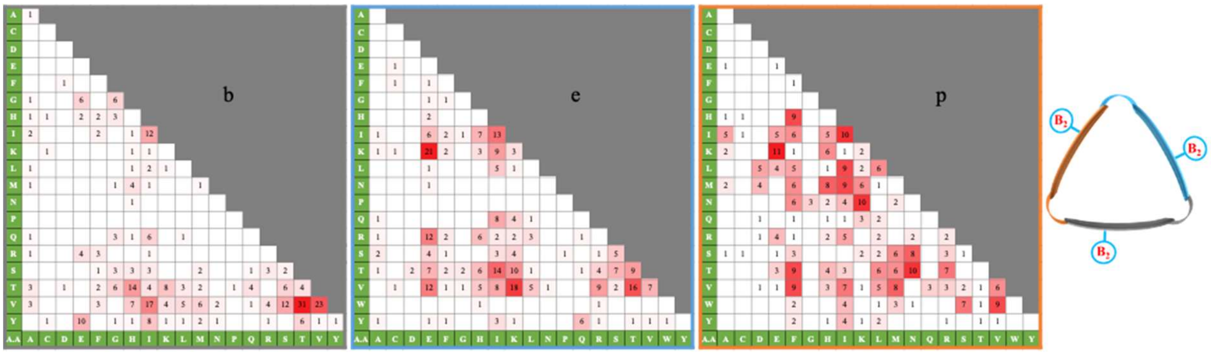


**Figure S9.** Rung-to-rung succession of residues at the $B_2$ position, for each face of the helix. The higher the number (indicated by intense red), the more frequent the succession.

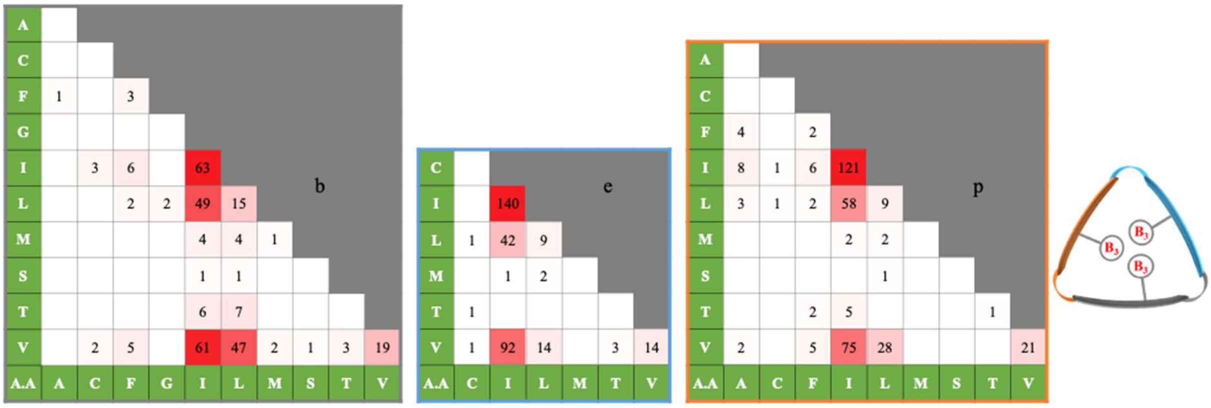


**Figure S10.** Rung-to-rung succession of residues at the $B_3$ position, for each face of the helix. The higher the number (indicated by intense red), the more frequent the succession.

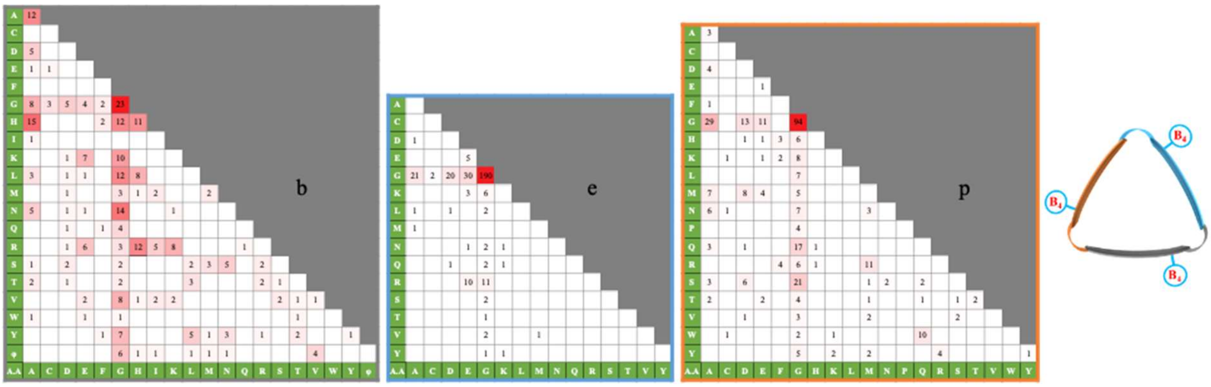


**Figure S11.** Rung-to-rung succession of residues at the $B_4$ position, for each face of the helix. The higher the number (indicated by intense red), the more frequent the succession.

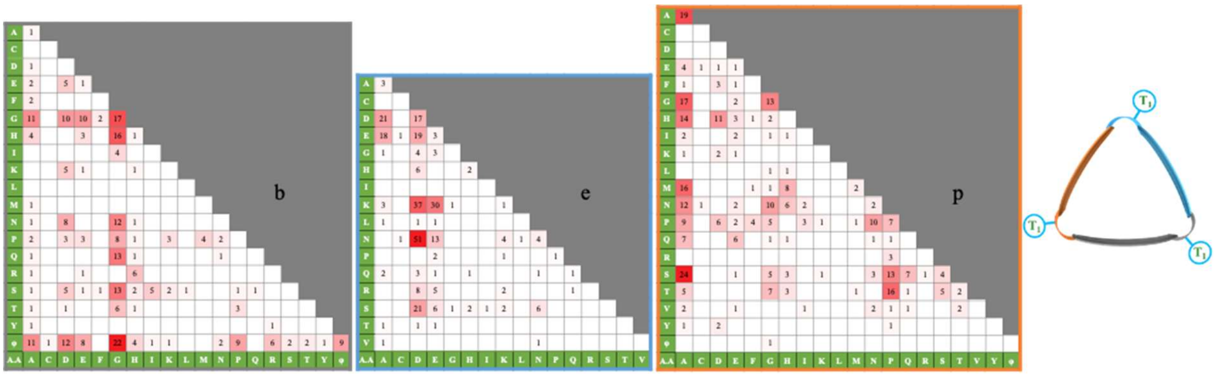


**Figure S12.** Rung-to-rung succession of residues at the $T_1$ position, for each face of the helix. The higher the number (indicated by intense red), the more frequent the succession.

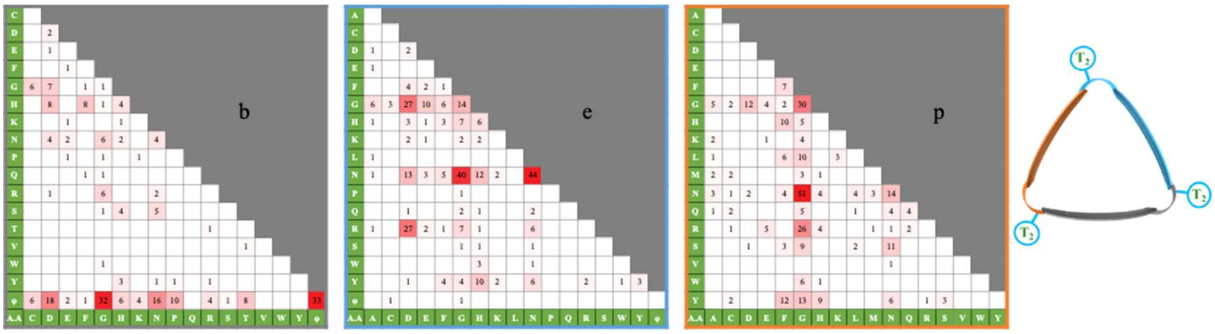


**Figure S13.** Rung-to-rung succession of residues at the $T_2$ position, for each face of the helix. The higher the number (indicated by intense red), the more frequent the succession.

The following figures (from S14 to S37) were generated the following way, using the Excel table in which all the protein sequences have been entered into an 18-rows table, with one line per rung. In order to analyze the interdependence of residues at key positions of a same rung, the function "UNIQUE" as well as the function "TRANSPOSE(UNIQUE) were used to create a matrix bearing 2 different positions. Then the "COUNTIF" function has been used with first argument the column of interest for the first position linked to the residue in the matrix (in column) and as second argument the one for the second position of interest linked to the second residue in the matrix (in line). This result of a matrix with interdependence of residues as a function of different positions. The matrix has then been colored based on the number of interdependences.

As an example, the resulting Figures can be read as follow: "Figure S14 shows that we observed 12 times the co-occurrence of an Ile residue at position $B_2$(N.b) and a Pro at position $T_1$(N.p)."

| T1 / B2 | P | S | H | T | D | V | E | φ | I | A | K | G | Q |
|---|---|---|---|---|---|---|---|---|---|---|---|---|---|
| I | 12 | 1 | | | | | 1 | | | | 1 | | |
| N | | 1 | | | | | | | | | | | |
| D | | | 1 | | | | | 12 | 1 | | 1 | | |
| R | 3 | 1 | | | | | | | 1 | | | 1 | 1 |
| V | 5 | 1 | | 2 | | | | | | 1 | | | |
| M | 2 | | | | 2 | | | | | | | | |
| Y | 3 | | | | | 1 | | 1 | | 1 | | | |
| K | | | | | 1 | | | | | | | | |
| A | 2 | | | | | | | | | | 1 | | |
| φ | | 1 | | | | | | 3 | | | | 4 | |
| F | | | | | | | | 1 | | | | | |
| L | 1 | | | | | | | | | | 1 | | |
| W | | | | | | | | 1 | | | | | |
| T | 2 | | | | | | | | | | | | |
| S | | | | | | | | | | | 1 | | |

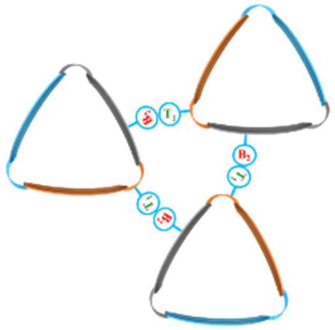

**Figure S14.** Co-occurrence of residues at positions $B_2$(N.b) and $T_1$(N.p), two positions facing each other within the trimer. ϕ symbol represents residues that are part of a loop.

| T1 / B2 | P | D | H | M | S | Q | E | I | N | G | V | A | F | T | Y | φ | K | R | L | C |
|---|---|---|---|---|---|---|---|---|---|---|---|---|---|---|---|---|---|---|---|---|
| S | 2 | 2 | 1 | | | | 6 | | 1 | 2 | 2 | 5 | | | | | | | | |
| V | 16 | 9 | 12 | 2 | 7 | 11 | 6 | | 1 | 2 | 1 | 19 | | 3 | | | | | 1 | |
| Q | 2 | | | 1 | | | | | | 3 | | 5 | | | | | | | | |
| A | 1 | | | | 1 | | | | | 5 | 1 | 1 | | 2 | | | | | | |
| I | 2 | 1 | 2 | 7 | 13 | 1 | 1 | | 1 | 3 | | 14 | | 4 | | | | | | |
| T | 6 | | 1 | 2 | 6 | 1 | 1 | 2 | 4 | 11 | | 16 | 3 | 2 | | | 1 | 1 | | 1 |
| H | 1 | 1 | 1 | | 1 | 1 | | | 12 | 2 | | | | 2 | | | | 1 | | |
| G | | | 1 | | 1 | | | | | 10 | | 9 | | | | | | | | |
| M | 2 | | 1 | 1 | 3 | | | 1 | | | | 4 | | | | | 1 | | | |
| L | | | | 1 | 1 | | | 1 | 1 | 2 | | 1 | 1 | | | | | | | |
| K | | | | | 2 | 1 | | 1 | | | | 7 | | 1 | | | | | | |
| F | 1 | | | | | | 1 | | 1 | 1 | | 1 | | 1 | | | 1 | | | |
| C | | | | | | | | 1 | | 1 | | 1 | | | | | | | | |
| Y | 3 | | | | | 1 | | | 2 | 1 | | 11 | | 5 | | | | | | |
| E | | | 8 | 6 | | | | | | | | | | | | | | | | |
| D | | | | | 1 | | | | | | | | | | | | | | | |
| R | | | | | 1 | | | | 4 | 3 | | 1 | | | | | | | | |
| N | 1 | | | | | | | | | 1 | | | | | | | | | | |
| P | | | | | | | | | | 1 | | | | | | | | | | |

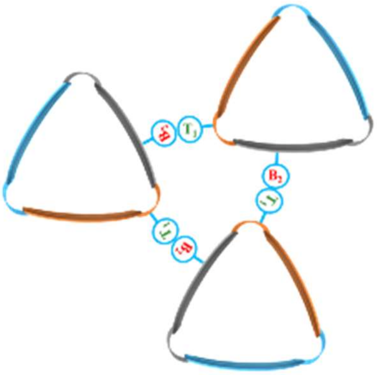

**Figure S15.** Co-occurrence of residues at positions $B_2$(I.b) and $T_1$(I.p), two positions facing each other within the trimer. ϕ symbol represents residues that are part of a loop.

| T1 / B2 | A | φ | S | G | Ø | V | K | T |
|---|---|---|---|---|---|---|---|---|
| V | 2 | 26 | 2 | | | 1 | | |
| G | | 4 | | | | | | |
| L | | 11 | | 1 | | | | |
| M | | 1 | 2 | 1 | | | | |
| S | | | | | 1 | | | |
| T | | 4 | 1 | | | | | 1 |
| W | | 2 | | | | | | |
| A | 1 | 4 | | | | | | |
| Y | 1 | 6 | | | | | | |
| R | | 1 | | | | | 1 | |
| F | | 1 | | | | | | |
| I | | 1 | | | | | | |
| K | | 1 | | | | | | |

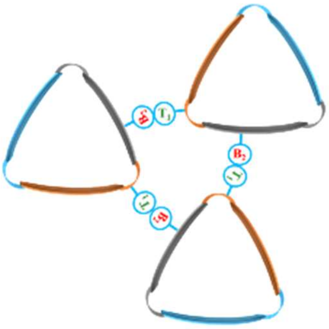

**Figure S16.** Co-occurrence of residues at positions $B_2$(C.b) and $T_1$(C.p), two positions facing each other within the trimer. ϕ symbol represents residues that are part of a loop.

| T2 / B2 | Q | T | F | φ | N | S | Y | G | P | R | L | D | K | H |
|---|---|---|---|---|---|---|---|---|---|---|---|---|---|---|
| I | 1 | 8 | | | 1 | 1 | | 2 | 1 | 1 | | | | |
| N | | | 1 | | | | | | | | | | | |
| D | | | | 13 | | 1 | | | 1 | | | | | |
| R | | 2 | | | 3 | | | 1 | | | | 1 | | |
| V | | 1 | | | 2 | 4 | | | 1 | | 1 | | | |
| M | | | | | | | 2 | | | 1 | | | | 1 |
| Y | | | 1 | 1 | 1 | | | 1 | 1 | 1 | | | | |
| K | | | | | 1 | | | | | | | | | |
| A | 1 | | | | | | | | 1 | | 1 | | | |
| φ | | | | 7 | | | | | | | | | 1 | |
| F | | | | 1 | | | | | | | | | | |
| L | | | | | | 1 | | 1 | | | | | | |
| W | | | | 1 | | | | | | | | | | |
| T | | | | | | | | | 1 | | | 1 | | |
| S | | | | | | | | | | | | | 1 | |

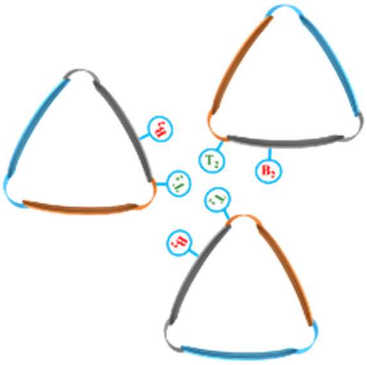

**Figure S17.** Co-occurrence of residues at positions $B_2$(N.b) and $T_2$(N.p). ϕ symbol represents residues that are part of a loop.. ϕ symbol represents residues that are part of a loop.

| T2 / B2 | M | G | Q | R | F | H | S | N | C | Y | A | V | L | K | E | φ | D | W |
|---|---|---|---|---|---|---|---|---|---|---|---|---|---|---|---|---|---|---|
| S | 1 | 4 | | | 1 | 1 | 4 | 7 | 1 | | | | 1 | | 1 | | | |
| V | 2 | 28 | 3 | 6 | 10 | 5 | 7 | 14 | 2 | 5 | 2 | | 1 | 2 | 1 | | 2 | |
| Q | 1 | 5 | 2 | | | | | 1 | | | | | | 1 | | | 1 | |
| A | | 3 | | 2 | | | | | 2 | 1 | | | 1 | 1 | 1 | | | |
| I | 1 | 23 | 1 | 2 | 2 | 8 | | 4 | | 2 | 1 | | | | 1 | | 3 | |
| T | 2 | 27 | 3 | | 1 | 2 | 2 | 9 | | 5 | | | 4 | | 1 | | 1 | |
| H | | 6 | | 2 | 3 | | 1 | 10 | | | | | | | | | | |
| G | 1 | 3 | 3 | 6 | | 1 | | 2 | | | 5 | | | | | | | |
| M | | 2 | | 4 | | | 1 | 1 | 1 | 3 | | | | 1 | | | | |
| L | | 4 | | 1 | | | | 1 | | | 1 | | | | | | | |
| K | | 7 | | | | | | 4 | | 1 | | | | | | | | |
| F | | 1 | | | | 1 | | 2 | | | | | | | | | | 1 |
| C | | 2 | | | | | | | | 1 | | | | | | | | |
| Y | | 9 | | | 1 | | 2 | 3 | | | | | | | | | | 5 |
| E | | 6 | | | 1 | | | | | | | | 7 | | | | | |
| D | | | | | | | | 1 | | | | | | | | | | |
| R | | 4 | | | 4 | | | | | 1 | | | | | | | | |
| N | | 1 | | | | | | 1 | | | | | | | | | | |
| P | | | | | | | | | | | 1 | | | | | | | |

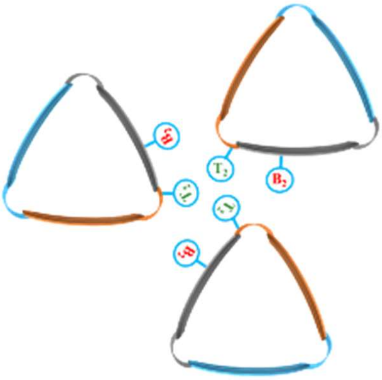

**Figure S18.** Co-occurrence of residues at positions $B_2$(I.b) and $T_2$(I.p). ϕ symbol represents residues that are part of a loop.

| B2 \ T2 | G | φ | M | Ø | R | N |
|---|---|---|---|---|---|---|
| V | 3 | 28 | | | | |
| G | | 4 | | | | |
| L | | 11 | | | 1 | |
| M | | 3 | 1 | | | |
| S | | | | 1 | | |
| T | | 5 | | | | |
| W | | 2 | | | | |
| A | 1 | 4 | | | | |
| Y | 1 | 6 | | | | |
| R | | 1 | | | | 1 |
| F | | 1 | | | | |
| I | | 1 | | | | |
| K | | 1 | | | | |

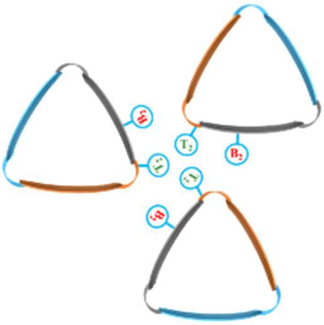

**Figure S19.** Co-occurrence of residues at positions $B_2$(C.b) and $T_2$(C.p). ϕ represents residues that are part of the C-ter capping motif (loop) and Ø missing residues.

| T1 \ T2 | Q | T | F | φ | N | S | Y | G | P | R | L | D | K | H |
|---|---|---|---|---|---|---|---|---|---|---|---|---|---|---|
| P | 2 | 8 | 1 | | 2 | 4 | | 3 | 5 | 2 | | 3 | | 1 |
| S | | 2 | 1 | | | | | | | | 1 | | 1 | |
| H | | | | 1 | | | | | | | | | | |
| T | | | | | 1 | 1 | | | | | | | | |
| D | | | | | 1 | 1 | 2 | | | | | | | |
| V | | | | | 1 | | | | | | | | | |
| E | | | | | | | | 1 | | | | | | |
| φ | | | | 18 | | | | | | | | | | |
| I | | | | | | | | 1 | 1 | | | | | |
| A | | | | | 1 | | | | | 1 | | | | |
| K | | 1 | | | | 2 | | | | | 1 | | 1 | |
| G | | | | 4 | 1 | | | | | | | | | |
| Q | | | | | 1 | | | | | | | | | |

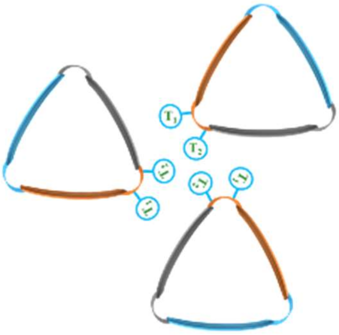

**Figure S20.** Co-occurrence of residues at positions $T_1$(N.p) and $T_2$(N.p). ϕ symbol represents residues that are part of a loop.

| T2 / T1 | M | G | Q | R | F | H | S | N | C | Y | A | V | L | K | E | φ | D | W |
|---|---|---|---|---|---|---|---|---|---|---|---|---|---|---|---|---|---|---|
| P | 1 | 12 | | 3 | 10 | 2 | 3 | 5 | | 12 | 1 | | 2 | | | | 1 | 1 |
| D | | 6 | 1 | | | | | 5 | 1 | | | | | | | | | |
| H | 1 | 2 | 3 | 4 | 1 | 2 | 1 | 1 | | 2 | 1 | 1 | 7 | 3 | 3 | | | |
| M | | 18 | 1 | | | 1 | | 2 | | | | | | | | | | |
| S | | 12 | | 4 | | 8 | 1 | 8 | | 2 | | | | | 1 | | 4 | |
| Q | | 4 | | | 6 | | 4 | 1 | | 1 | | | | | | | 1 | |
| E | | 4 | 1 | | 5 | 1 | 2 | 1 | 1 | 1 | | | | | | | | |
| I | | | | | | | 1 | 4 | | 1 | | | | | | | | |
| N | | 4 | 2 | | 6 | | 1 | 9 | | | | | 2 | | | | | |
| G | 5 | 17 | 2 | 1 | | 2 | 1 | 3 | 2 | 1 | 7 | | 3 | 1 | 1 | | | |
| V | | 2 | | | | | | 4 | | | | | | | | | | |
| A | 1 | 53 | 2 | 9 | 1 | 2 | 2 | 19 | 2 | 8 | | | | 2 | | | | |
| F | | 2 | | 1 | | | | 5 | | | | | | | | | 2 | |
| T | | 9 | | | | | 1 | 3 | | | 1 | | 1 | | 3 | | | 6 |
| Y | | 2 | | | | | | 1 | | | | | | | | | 1 | |
| φ | | | | | | | | | | | | | | | | 1 | | |
| K | | | | | | | | 1 | | | | | | | | | 1 | |
| R | | | | 1 | | | | | | | | | | | | | | |
| L | | 1 | | | | | | | | | | | | | | | | |
| C | | | | | | | | | | 1 | | | | | | | | |

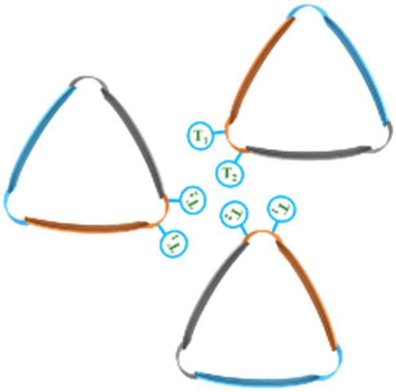

**Figure S21.** Co-occurrence of residues at positions $T_1$(I.p) and $T_2$(I.p). ϕ symbol represents residues that are part of a loop.

| T2 / T1 | G | φ | M | Ø | R | N |
|---|---|---|---|---|---|---|
| A | 4 | | | | | |
| φ | | 62 | | | | |
| S | 1 | 3 | | | | |
| G | | | 1 | | 1 | |
| Ø | | | | 1 | | |
| V | | 1 | | | | |
| K | | | | | | 1 |
| T | | 1 | | | | |

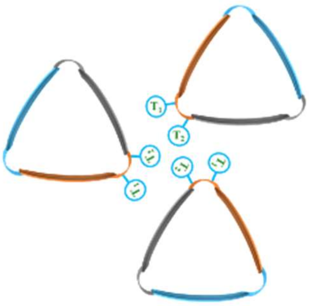

**Figure S22.** Co-occurrence of residues at positions $T_1$(C.p) and $T_2$(C.p). ϕ represents residues that are part of the C-ter capping motif (loop) and Ø missing residues.

| T1 \ T2 | T | S | D | G | H | R | E | N | φ | Y | P | Q |
|---|---|---|---|---|---|---|---|---|---|---|---|---|
| P | 3 | 2 | | | | | 1 | 1 | | | | |
| K | | 3 | | 3 | 1 | 1 | | 5 | | 1 | | 1 |
| S | 1 | 1 | 1 | 1 | | 1 | | | | | | |
| H | | | 1 | 1 | 1 | | | | | | | |
| A | | 1 | 1 | 1 | 1 | | | | | | | |
| E | 3 | 2 | 1 | 3 | | 1 | | 2 | | 1 | | |
| Q | | | | 1 | | 2 | | | | | | |
| D | | 2 | 1 | 2 | 1 | 1 | | 2 | | 1 | | 1 |
| G | | | | | 1 | | | | | | | |
| R | | 1 | 2 | 3 | | | | 2 | | 1 | | |
| φ | | | | | | | | | 2 | | | |
| V | | | | | | | 1 | | | | | |
| T | | | | | | | 1 | | | | | |
| M | | | | | | | | | | | 1 | |
| C | | | | 1 | | | | | | | | |
| N | | | | | | | | 1 | | | | |

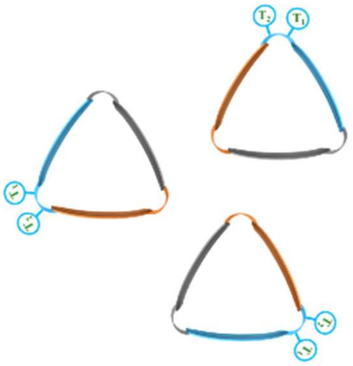


**Figure S23.** Co-occurrence of residues at positions $T_1$(N.e) and $T_2$(N.e). ϕ symbol represents residues that are part of a loop.

| T1 \ T2 | N | R | D | F | H | Y | E | Q | W | K | A | P | L | C | φ | S |
|---|---|---|---|---|---|---|---|---|---|---|---|---|---|---|---|---|
| A | 4 | 4 | 3 | 1 | 4 | 1 | 4 | 2 | | | 2 | | | | | |
| D | 39 | 11 | 24 | 5 | 13 | 8 | 1 | | 2 | 2 | 2 | | 1 | | | |
| N | 21 | 4 | 9 | 1 | 1 | 2 | 2 | 1 | | 1 | | | | | | |
| E | 12 | 1 | 6 | | 1 | 3 | 3 | | 1 | 3 | | | | 1 | | 1 |
| R | 2 | | 2 | 1 | 1 | 2 | | | | | | | | | | |
| S | 6 | 1 | 1 | 3 | 6 | 4 | | | | | 1 | | | 1 | | |
| Q | 2 | | | | 1 | | | | | | | | | 1 | | |
| K | 18 | 3 | 3 | 5 | 4 | 2 | | | | | 1 | | | | | 1 |
| L | | | | 1 | | | | | | | | | | | | |
| G | | 1 | | | 1 | 1 | | | | | 3 | 1 | | | 1 | |
| H | 3 | 1 | 1 | | | | 1 | | | | | | | | | |
| I | | | | | | | | | | | | | | | | |
| P | | | | | | | 1 | | | 1 | | | | | | |
| T | 1 | | | | | | | | | | | | | | | |
| C | 1 | | | | | | | | | | | | | | | |
| V | | | | | 1 | | | | | | | | | | | |

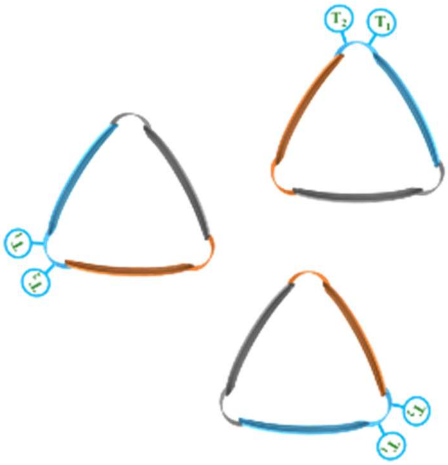


**Figure S24.** Co-occurrence of residues at positions $T_1$(I.e) and $T_2$(I.e). ϕ symbol represents residues that are part of a loop.

| T2 / T1 | G | Y | P | Ø | F | N | D | E | R | H | S | φ | Q | W |
|---|---|---|---|---|---|---|---|---|---|---|---|---|---|---|
| D | 2 | 1 | | | 1 | 5 | | | 1 | | 1 | | | 1 |
| P | 2 | 12 | | | 3 | 4 | | | 1 | 4 | | | | |
| A | 5 | 3 | | 1 | | 1 | 2 | | | 2 | | | | |
| E | 1 | 1 | 2 | | | 3 | | | | | | | 1 | |
| G | | | | 1 | | 1 | | | | | 1 | | | |
| S | 3 | | 1 | | | 2 | | | | | | | | |
| K | | 1 | | 1 | | | | 1 | | | | | | |
| Y | 1 | | | | | | | | | | | | | |
| Q | | | | | | | | | | | | 1 | | |
| N | 2 | | | | | 1 | | | | | | | | |
| H | | | | | | 1 | | | | | | | | |

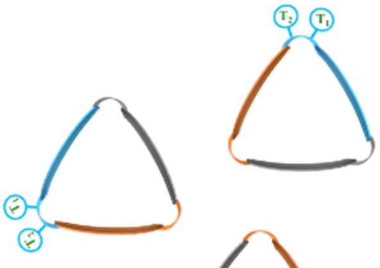

**Figure S25.** Co-occurrence of residues at positions $T_1$(C.e) and $T_2$(C.e). ϕ represents residues that are part of the C-ter capping motif (loop) and Ø missing residues.

| T2 / T1 | φ | G | S | D | A | E | N | H | K | T | P |
|---|---|---|---|---|---|---|---|---|---|---|---|
| φ | 17 | | | | | | | | | | |
| E | | 4 | 1 | 1 | | | 1 | | 1 | | |
| P | | 3 | 6 | | 7 | 1 | 1 | | 2 | 1 | 1 |
| A | 1 | | | 1 | | | | | | 1 | |
| G | | | | 3 | 1 | 1 | 1 | | 1 | 3 | |
| D | | | | | | 1 | | 1 | | | |
| I | | | | 2 | | | | | | | |
| L | | 3 | | | | | | | | | |
| F | | | | | | | 1 | | | | |
| Q | 1 | 2 | | | | | | | | | |
| H | 1 | | 1 | | | | | | | | |
| V | | | | 1 | | | | | | | |
| K | | | 1 | | | | | | | | |
| R | | 3 | | | | | | | | | |

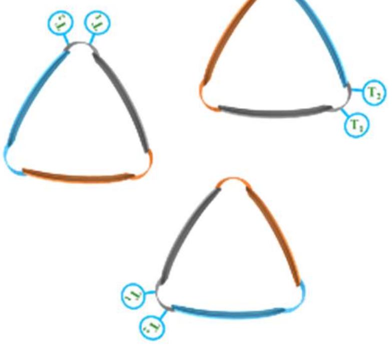

**Figure S26.** Co-occurrence of residues at positions $T_1$(N.b) and $T_2$(N.b). ϕ symbol represents residues that are part of a loop.

| T2 \ T1 | E | φ | P | K | H | D | F | Q | G | R | N | S | V | T | Y | C | W |
|---|---|---|---|---|---|---|---|---|---|---|---|---|---|---|---|---|---|
| G | 2 | 25 | 5 | 1 | 19 | 7 | 1 | 2 | 4 | 4 | 15 | 5 | | | | | |
| S | | 2 | | 1 | 1 | 1 | | | 2 | 1 | 2 | 1 | | | | 6 | |
| φ | | 53 | | | | | | | | | | | | | | | |
| F | | | | 1 | | | | | | | 3 | | | | | | |
| P | | 2 | | | 1 | | | 1 | 19 | | 4 | | | | 1 | | |
| R | | 6 | | | | | | | | | 2 | | | | | | |
| H | 2 | | | | 1 | 14 | | | 2 | | 2 | | | | | | |
| Q | | 2 | | | 1 | | 10 | 1 | | | | | | | 2 | | |
| A | 1 | 4 | 7 | 1 | | 1 | | | 8 | 2 | 1 | | | 1 | 2 | | |
| N | | | | 1 | | 1 | | | 3 | 1 | | | | | | | |
| D | | 1 | | | 2 | | | | 7 | 2 | 5 | | 1 | | | | |
| E | | | | 1 | | 1 | | | 8 | 1 | 3 | | | 8 | | | |
| C | | | | | | | | | 1 | | | | | | | | |
| T | | 6 | | | | | | | | | | | | | | | 1 |
| I | | | | | | | | | 6 | | | | | | | | |
| K | | 1 | | | | 1 | | | | | 3 | | | | | | |
| M | | 4 | | | | | | | 1 | | | | | | | | |
| L | | | | | | | | | 1 | | | | | | | | |
| Y | | | | | | | | | 1 | | 1 | | | | | | |

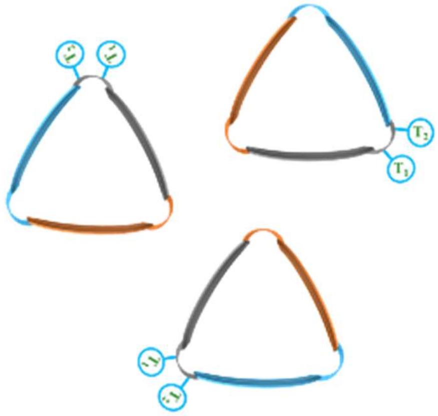

**Figure S27.** Co-occurrence of residues at positions $T_1$(I.b) and $T_2$(I.b). ϕ symbol represents residues that are part of a loop.

| T2 \ T1 | Ø | G | H | N | K | R | φ |
|---|---|---|---|---|---|---|---|
| Q | 12 | 1 | 1 | 1 | | | 1 |
| K | 27 | | | | | | |
| R | 8 | 1 | | | | | |
| N | 1 | | | | | | |
| A | | 1 | | | | | |
| P | | 2 | | 1 | | | |
| E | 4 | 1 | | | 1 | | |
| I | 1 | | | | | | |
| T | | | | | | | |
| G | 1 | | | | | | |
| S | | | | 1 | | | |
| V | | | | | | 1 | |
| C | 1 | | | | | | |
| φ | | | | | | | 4 |

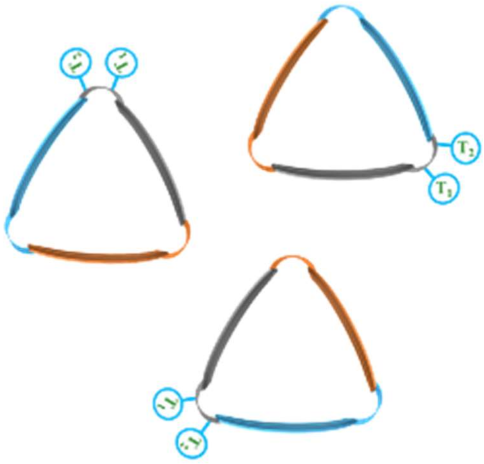

**Figure S28.** Co-occurrence of residues at positions $T_1$(C.b) and $T_2$(C.b). ϕ represents residues that are part of the C-ter capping motif (loop) and Ø missing residues.

| T1 / B4 | φ | E | P | A | G | D | I | L | F | Q | H | V | K | R |
|---|---|---|---|---|---|---|---|---|---|---|---|---|---|---|
| E | 1 | 4 | 1 | | | | | | | | | | | |
| D | 5 | 1 | 3 | 1 | | | 2 | | | | | 1 | 1 | |
| A | | | | 1 | | | | | | | | | | |
| I | | | | | 4 | | | | | | | | | |
| H | | 1 | 13 | | | | | | | | | | | |
| L | | | | | | 1 | | | | | 1 | | | |
| Y | 4 | | | 1 | | | | | | | | | | |
| G | | 1 | | | | | | | | | | | | |
| R | | 1 | | | 4 | 1 | | 3 | | 2 | 1 | | | 3 |
| K | | | | | 1 | | | | | | | | | |
| N | | | | | | | | | 1 | | | | | |
| S | 1 | | 4 | | | | | | | | | | | |
| V | | | 1 | | 1 | | | | | 1 | | | | |
| φ | 5 | | | | | | | | | | | | | |
| Q | 1 | | | | | | | | | | | | | |

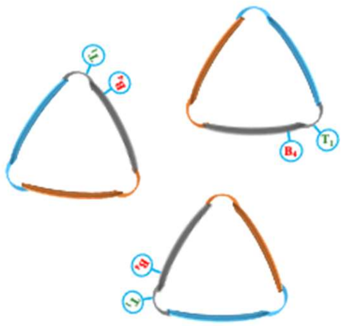

**Figure S29.** Co-occurrence of residues at positions $B_4$(N.b) and $T_2$(N.b). ϕ symbol represents residues that are part of a loop.

| T1 / B4 | P | K | S | H | A | E | Q | D | G | R | φ | V | T | M | C | N |
|---|---|---|---|---|---|---|---|---|---|---|---|---|---|---|---|---|
| D | 2 | 2 | 1 | | | 1 | | | | | | | | | | |
| A | 3 | 3 | 3 | | 3 | | | 4 | | 3 | 2 | | | | | |
| G | | 8 | 1 | 3 | 1 | 7 | 3 | 6 | | 6 | | | | | 1 | 1 |
| S | | 1 | | | | 1 | | | | | | | | | | |
| K | | | | | | | | | 1 | | | | | | | |
| E | | | | | | 1 | | 1 | | | | 1 | | | | |
| H | 2 | | | | | 3 | | | | | | | | | | |
| V | | | | | | | | | | | | | 1 | | | |
| L | | | | | | | | | | | | | | 1 | | |
| N | | 1 | | | | | | | | | | | | | | |

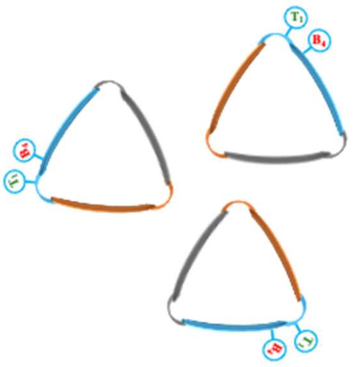

**Figure S30.** Co-occurrence of residues at positions $B_4$(N.e) and $T_2$(N.e). ϕ symbol represents residues that are part of a loop.

| T1 / B4 | P | S | H | T | D | V | E | φ | I | A | K | G | Q |
|---|---|---|---|---|---|---|---|---|---|---|---|---|---|
| D | 3 | 1 | 1 | 1 | 1 | | | 8 | | | | | |
| H | 13 | 3 | | | | | | | | | 2 | | |
| E | 5 | 1 | | | 1 | | | | | | | | 1 |
| S | | | | 1 | | | | | 1 | | 1 | 4 | |
| A | 3 | | | | 2 | | | 3 | | | 2 | 1 | |
| F | | | | | | 1 | | | | | | | |
| K | | | | | | | 1 | | | 1 | | | |
| L | | | | | | | | 2 | | | | | |
| R | | | | | | | | 1 | | | | | |
| Y | | | | | | | | 1 | | | | | |
| V | 6 | | | | | | | 2 | | | | | |
| N | 1 | | | | | | | | 1 | | | | |
| G | | | | | | | | 1 | | 1 | | | |

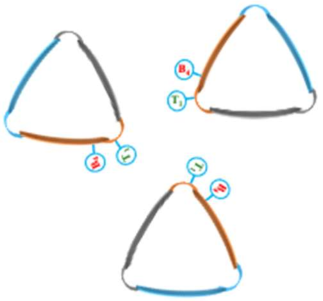

**Figure S31.** Co-occurrence of residues at positions $B_4$(N.p) and $T_2$(N.p). ϕ symbol represents residues that are part of a loop.

| T1 / B4 | G | S | φ | F | P | R | H | Q | A | N | D | E | C | T | I | K | M | L | Y |
|---|---|---|---|---|---|---|---|---|---|---|---|---|---|---|---|---|---|---|---|
| I | 7 | | | | | | | | 2 | | | | | | | | 2 | | |
| R | 8 | 1 | 1 | | 8 | | | 1 | 2 | | 1 | 2 | | | | | | | |
| H | 11 | 2 | 6 | | 1 | 3 | | 13 | 2 | | | | | 4 | | 1 | | | |
| V | 1 | | 3 | 1 | | | | | 5 | | 2 | 6 | 1 | | | | | | |
| G | 12 | 11 | 21 | | 2 | 1 | 2 | | 6 | 3 | 11 | 12 | | | 4 | 3 | | | |
| N | 5 | 1 | | 3 | 5 | 1 | | | 2 | | | | | | 6 | | | 1 | 1 |
| A | 18 | | | | | | 13 | | 3 | | | | | | | | | | |
| E | 7 | 2 | | | | | 1 | | | 1 | 3 | 2 | | | | | | | |
| Y | 2 | | 3 | | | | 2 | | 1 | | 1 | | | 2 | | | | | |
| D | 1 | | 2 | | | 3 | 2 | | | | 1 | | | 1 | | | | | |
| S | 2 | 2 | 1 | | | | | | 3 | 1 | | | | | | 1 | 1 | | |
| F | 1 | | | | 1 | | | | | 2 | | | | | | | | | |
| Q | 1 | | | | | | 1 | | 1 | | 1 | | | | | | | | |
| L | 11 | 1 | 4 | | 5 | | | | 1 | 2 | 6 | 2 | | | | | | | 1 |
| C | 1 | | | | | | | 1 | 1 | | | | | | | | | | |
| T | 4 | 1 | 3 | | 1 | | | | | | 1 | | | | | | | | |
| K | 5 | | | | | | | 1 | | 6 | | 1 | | | | 3 | | | |
| φ | | | 8 | | | | | | | | | | | | | | | | |
| M | 2 | | 1 | | 6 | | | | | | 2 | | | | | | 2 | | |
| W | | | | | | | | | | | 1 | 1 | | | | | | | |

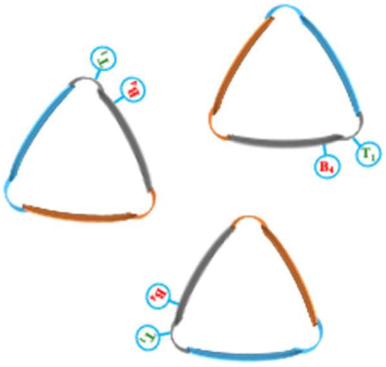

**Figure S32.** Co-occurrence of residues at positions $B_4$(I.b) and $T_2$(I.b). ϕ symbol represents residues that are part of a loop.

| T1 / B4 | A | D | N | E | R | S | Q | K | L | G | H | I | P | T | C | V |
|---|---|---|---|---|---|---|---|---|---|---|---|---|---|---|---|---|
| G | 30 | 79 | 43 | 44 | 11 | 23 | 4 | 42 | 1 | 4 | 8 | | 3 | 1 | 1 | 1 |
| D | 2 | 6 | | 6 | | 1 | | 1 | | | | | | | | |
| C | | 2 | | | | | | | | | | | | | | |
| K | | 2 | | | | | | 4 | | | | | | 1 | | |
| E | | 21 | 4 | 7 | | | | 2 | 1 | 1 | | | | | | |
| R | | 1 | | 7 | | | 2 | 1 | | | | | | | | |
| A | 3 | 7 | 3 | 2 | | | | 1 | | | | | | | | |
| N | | 1 | | 1 | | | | | | | | 1 | | | | |
| V | | | | 2 | | | | | | | | | | | | |
| M | | | | 1 | | | | | | | | | | | | |
| T | | | | | | | | | | 1 | | | | | | |
| L | | | | 1 | | | | | | 2 | | | | | | |
| Q | | 2 | | 1 | | | | | | | | | | | | |
| S | | 1 | | | | | | | | | | | | | | |
| Y | | 1 | | | | | | | | | | | | | | |

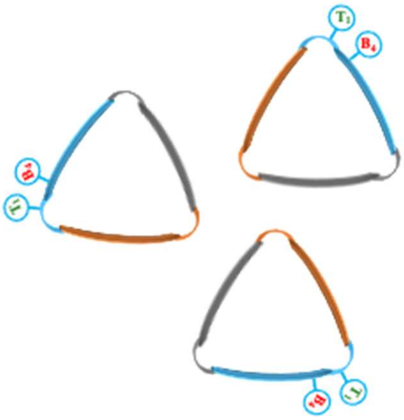

**Figure S33.** Co-occurrence of residues at positions $B_4$(I.e) and $T_2$(I.e). ϕ symbol represents residues that are part of a loop.

| T1 / B4 | P | D | H | M | S | Q | E | I | N | G | V | A | F | T | Y | φ | K | R | L | C |
|---|---|---|---|---|---|---|---|---|---|---|---|---|---|---|---|---|---|---|---|---|
| S | 4 | | | 5 | 3 | | | | | 7 | 1 | 1 | | | | | | | 1 | |
| Q | 1 | 10 | | | 3 | | 1 | | 1 | 1 | | 1 | | | | | | | | |
| A | 7 | | 2 | | 5 | | | | 8 | 7 | | 10 | | | | | | | | |
| G | 26 | | 18 | 12 | 17 | 1 | 1 | 3 | 7 | 26 | 1 | 73 | 1 | 10 | | | 3 | 1 | | |
| E | 2 | 2 | | | | | 1 | | | 3 | | 5 | | | | | | | | |
| K | 1 | 1 | | | 5 | 1 | | | | | | | | | | | | | | |
| Y | 2 | | | | 1 | 6 | | | | | | | | | | | | | | |
| R | | | | | 1 | | 11 | | | | | | | 1 | | | | | | |
| M | 7 | | 3 | 3 | | | 1 | 2 | | 2 | 2 | 8 | | 1 | | 1 | | | | 1 |
| H | 2 | | | | 3 | 1 | | | 1 | | | | | | | | | | | |
| F | 2 | | | | 1 | 2 | | | | | | | | | | | | | | |
| D | | | | 2 | 2 | | 1 | | 4 | | 2 | | | 8 | 1 | | | | | |
| T | | | 2 | | | | | | 4 | 2 | | | | | | | | | | |
| L | | | 2 | | | 5 | | | | | | | | | | | | | | |
| W | 4 | | 1 | | | 1 | | | | | | 1 | 4 | | 2 | | | | | |
| C | 1 | | | | | | | | | | | 1 | | | | | | | | |
| N | | | 2 | | | | | 1 | | | | | 5 | 4 | 1 | | | | | |
| P | | | 2 | | | | | | | | | | | | | | | | | |
| V | | | | | | | | | 2 | | | 1 | | | | | | 1 | | |

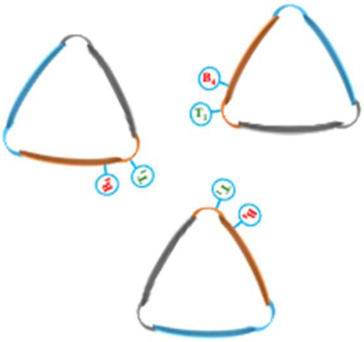

**Figure S34.** Co-occurrence of residues at positions $B_4$(I.p) and $T_2$(I.p). ϕ symbol represents residues that are part of a loop.

| T1 / B4 | Q | K | R | N | A | P | E | I | T | G | S | V | C | φ |
|---|---|---|---|---|---|---|---|---|---|---|---|---|---|---|
| T | | 16 | 5 | | | | 2 | | 1 | 1 | 1 | | | 2 |
| A | 2 | | | | | | | | | | | | | |
| L | 4 | 2 | | 1 | | | 1 | 1 | | | | | | |
| N | | 1 | 1 | | | | | | | | | | | |
| P | 2 | | | | 1 | 3 | | | | | | | | |
| D | | | 1 | | | | | | | | | | | |
| V | 2 | 3 | 1 | | | | 1 | | | | | 1 | 1 | |
| K | | | | | | | 2 | | | | | | | |
| S | 3 | 1 | | | | | | | | | | | | |
| G | 3 | 4 | | | | | | | | | | | | 2 |
| M | | | 1 | | | | | | | | | | | |

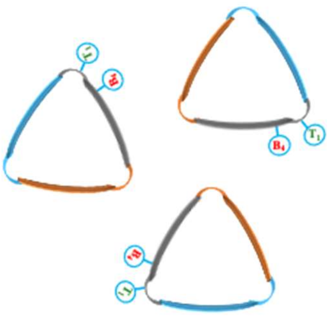


**Figure S35.** Co-occurrence of residues at positions $B_4$(C.b) and $T_2$(C.b). ϕ represents residues that are part of the C-ter capping motif (loop) and Ø missing residues.

| T1 / B4 | D | P | A | E | G | S | K | Y | Q | N | H |
|---|---|---|---|---|---|---|---|---|---|---|---|
| P | 8 | 20 | 12 | 2 | 1 | | 2 | | | 2 | 1 |
| T | | | | 2 | | 1 | | | | | |
| E | | 2 | 1 | | | 2 | 1 | | | | |
| V | | 1 | | | 1 | | | | | | |
| D | 1 | | | | | 1 | | | | | |
| G | 3 | | | 2 | | | | 1 | 1 | | |
| A | | 3 | 1 | 2 | 1 | | | | | | |
| K | | | | | | 1 | | | | | |
| S | | | | | | 1 | | | | 1 | |

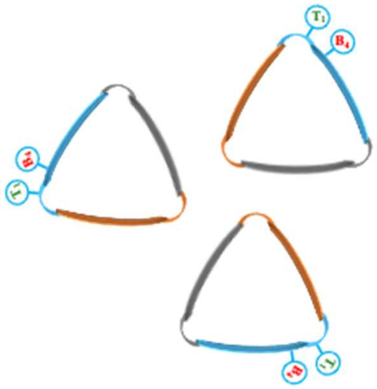


**Figure S36.** Co-occurrence of residues at positions $B_4$(C.e) and $T_2$(C.e). ϕ represents residues that are part of the C-ter capping motif (loop) and Ø missing residues.

| T1 / B4 | A | φ | S | G | Ø | V | K | T |
|---|---|---|---|---|---|---|---|---|
| P | 2 | | 1 | | | | | |
| Q | | 2 | | | | | | |
| S | | 3 | 2 | | | | | |
| A | | 13 | | | | | | |
| V | | 16 | | | | | | |
| G | | 8 | 1 | 1 | | | | |
| T | | 1 | | | 1 | | | |
| L | | 4 | | 1 | | | | |
| I | | 5 | 1 | | | | | 1 |
| Y | | 1 | | | | | | |
| K | | | | | | 1 | | |
| D | 1 | 1 | | | | | | |
| E | 1 | 2 | | | | | 1 | |
| F | | 3 | | | | | | |
| M | | 1 | | | | | | |
| R | | 2 | | | | | | |

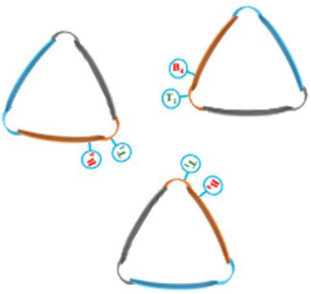

**Figure S37.** Co-occurrence of residues at positions $B_4$(C.p) and $T_2$(C.p). ϕ represents residues that are part of the C-ter capping motif (loop) and Ø missing residues.

| $B_4$ | | $T_1$ | $T_2$ | |
|---|---|---|---|---|
| H | 13 | Pro | A | 7 |
| S | 4 | | S | 6 |
| D | 3 | | A | 7 |
| E | 1 | | G | 3 |
| V | 1 | | K | 2 |
| | | | E | 1 |
| | | | P | 1 |
| | | | T | 1 |

| $B_4$ | | $T_1$ | $T_2$ | |
|---|---|---|---|---|
| A | 3 | Pro | T | 3 |
| D | 2 | | S | 2 |
| H | 2 | | E | 1 |
| | | | N | 1 |

| $B_4$ | | $T_1$ | $T_2$ | |
|---|---|---|---|---|
| H | 13 | Pro | T | 8 |
| V | 6 | | P | 5 |
| E | 5 | | S | 4 |
| A | 3 | | D | 3 |
| D | 3 | | G | 3 |
| N | 1 | | N | 2 |
| | | | Q | 2 |
| | | | R | 2 |
| | | | F | 1 |
| | | | H | 1 |

**Figure S38.** Occurrence of residues at positions $B_4$ and $T_2$, when a Pro occupies the $T_1$ position, at the N-ter rung, for the different hexads (from left to right: b, e and p).

| $B_4$ | | $T_1$ | $T_2$ | |
|---|---|---|---|---|
| P | 20 | Pro | Y | 12 |
| A | 3 | | H | 4 |
| E | 2 | | N | 4 |
| V | 1 | | F | 3 |
| | | | G | 2 |
| | | | R | 1 |

**Figure S39.** Occurrence of residues at positions $B_4$(C.e) and $T_2$(C.e), when a Pro occupies the $T_1$(C.e) position. It shows the frequency of the motif Pro-Pro-Tyr.

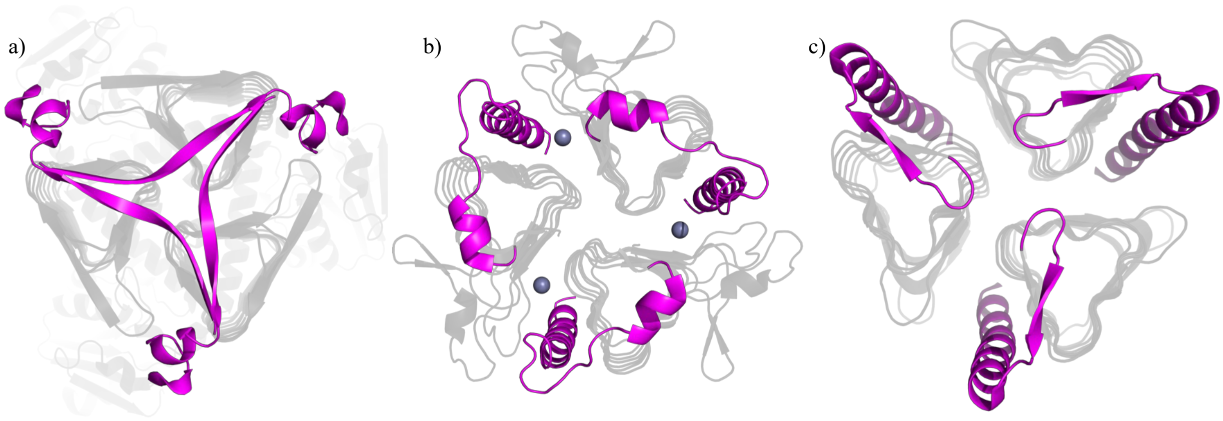


**Figure S40.** Examples capping motifs (in pink) that participate to the interaction of the other monomers within the homo-trimer of LβH-I. a) Long β strand crossing the whole trimer covering the hydrophobic core of another helix (PDB code 1HM9). b) α-helix covering the hydrophobic core of "its" helix and then form a α-helix along the (p) hexad in a groove at the interface of two monomers of LβH-I (PDB code 3OTM). c) β-hairpin motif (most commonly found) and α-helix along the (p) hexad in the groove between two monomers of LβH-I (PDB code 1XHD).
